# PDK1 and YAP1/TEAD signaling drive acquired KRAS inhibitor resistance in KRAS-mutant non-small cell lung cancer

**DOI:** 10.64898/2025.12.22.696060

**Authors:** Amrita Barua, Samrat T. Kundu, Sofia G. Garza, Jared J. Fradette, Bhagya Shree Choudhary, David H. Peng, Lixia Diao, Jing Wang, Don L. Gibbons

**Author notes:** Corresponding authors: Don L. Gibbons,; Samrat T. Kundu,. Contributed equally to this work (Amrita Barua, Samrat T. Kundu).

## Abstract

Mutations in KRAS are responsible for driving approximately 30% of NSCLC. While historically considered undruggable, recent breakthroughs have seen the FDA approval of two potent KRAS^G12C^ inhibitors, sotorasib (AMG510) and adagrasib (MRTX849). However, the efficacy of these inhibitors in the clinics has been limited by primary and acquired means of resistance. To elucidate mechanisms of acquired resistance, we generated a panel of resistant cell lines to the allele-specific KRAS inhibitors MRTX849 and MRTX1133 and observed an increased activation of the PDK1 and YAP1/TEAD signaling pathways. Pharmacological inhibition and genetic loss-of-function studies revealed a strong dependence on these pathways for the generation and maintenance of resistance to KRAS inhibition, which was then validated *in vitro* and *in vivo*. Furthermore, overexpression studies revealed that forced expression of either PDK1 or YAP1 led to increased resistance to KRAS inhibition in the sensitive lines. Taken together, our findings suggest that co-targeting PDK1 or YAP1/TEAD might be a potential approach to overcoming resistance to KRAS inhibition in NSCLC.

## Introduction

The Kirsten rat sarcoma viral oncogene homolog (*KRAS*) is one of the most frequently mutated oncogenic drivers and contributes to the development of multiple lethal cancers, including about 86% of pancreatic adenocarcinoma (PDAC), 41% of colorectal cancer (CRC), and 32% of non-small cell lung cancer (NSCLC) ^1,2^. KRAS is a GTPase located at a critical signaling junction downstream of various receptor tyrosine kinases (RTKs), such as the epidermal growth factor receptor (EGFR) family of receptors. It acts as a cellular switch by toggling between an active GTP-bound “on” state and a GDP-bound “off” state. Upon activation, KRAS transduces the receptor-mediated external stimuli into downstream signaling cascades, such as the mitogen-activated protein kinase (MAPK) and phosphoinositide 3-kinase (PI3K) pathways ^2,3^. These pathways are essential to the cell because they promote cell growth and survival processes. The oncogenic mutations in KRAS occur predominantly in codons 12, 13, and 61, and lock the protein in a prolonged active state which leads to dysregulation of these pathways and uncontrolled cell proliferation.

Until a few years ago, direct inhibition of KRAS was therapeutically challenging due to a lack of obvious drug-binding pockets in the structure of the protein. However, after decades of research, in 2013, while developing molecular inhibitors that selectively target KRAS^G12C^, Ostrem and colleagues discovered a previously unknown groove within the switch II region of the KRAS protein ^4^. This seminal finding led to the discovery and testing of multiple KRAS inhibitors, ultimately resulting in the FDA approval of two KRAS^G12C^ -specific inhibitors, AMG510 (sotorasib) in 2021 and MRTX849 (adagrasib) in 2022 ^5–12^.

While these drugs clearly demonstrate clinical efficacy for NSCLC patients, their benefits are modest with objective response rates (ORR) of about 40% and median progression-free survival (PFS) of about 6.5 months ^6,11^. The efficacy of these therapies has been restricted by primary and/or acquired means of resistance, including but not limited to activating mutations in other hotspots of KRAS, compensatory overactivation of WT KRAS along with other RAS family members, amplifications of RTKs such as MET and FGFR2, and histological transformation ^13,14^. Therefore, research is being carried out to develop second generation KRAS^G12C^ inhibitors with improved efficacy, along with compounds that target other mutations of KRAS, such as KRAS^G12D^, and pan-KRAS and pan-RAS inhibitors. Several such inhibitors are currently undergoing evaluation in preclinical and clinical settings, but none so far have been reported to result in dramatically increased response or PFS ^15^. Therefore, there remains a critical need for finding non-genomic molecular determinants of resistance in order to formulate combination treatment regimens which can delay the onset of resistance to KRAS inhibition in patients.

In this paper, we utilized the previously established 344SQ-G12D cell lines derived from the clinically relevant KRAS^G12D(LA1)^/p53^R172HΔG/+^ (KP) mouse model and used CRISPR/Cas9 to generate KRAS^G12C^-based *in vitro* models (344SQ-G12C) ^16,17^. We then subjected these lines to prolonged treatment with increasing doses of MRTX849 (G12Ci) or MRTX1133 (G12Di) until they acquired resistance to KRAS inhibition (KRASi). Interrogation of the resistant lines revealed heightened activation of 3-phosphoinositide-dependent kinase-1 (PDK1) and Yes-associated protein 1/TEA domain transcription factor (YAP1/TEAD) pathways in these lines compared to their sensitive counterparts. Pharmacological suppression revealed that co-targeting PDK1 or TEAD along with KRAS improved the sensitivity to KRASi in the resistant models *in vitro* and *in vivo*. Using genetic knockout models, we also demonstrated that loss of PDK1 or YAP1 is sufficient to resensitize the resistant cells to KRAS inhibition in cell lines and mouse models. Furthermore, using PDK1 and YAP1 overexpression cells, we illustrated that forced overexpression of these proteins is sufficient to impart resistance to KRAS inhibition *in vitro.* While the YAP1/TEAD pathway has been implicated in giving rise to KRASi resistance by multiple groups, our research indicates that PDK1 is also a driver of acquired resistance to KRASi in NSCLC ^18–20^. Surprisingly, we also find that knocking out or overexpressing PDK1 leads to alterations in YAP1 levels in the cells, suggesting that PDK1 might be a potential upstream regulator of YAP1. Overall, our findings demonstrate that targeting PDK1 and YAP1/TEAD pathways might be a possible strategy to overcome resistance to KRAS inhibition in NSCLC.

## Results

To understand the mechanisms of adaptive resistance to direct KRAS mutant allele-specific inhibitors (KRASi), we utilized both our established KP (*Kras^G12D-LA1/+^; TP53^R172HτιG/+^*) murine syngeneic cell line models, ^16,17^ and widely available human NSCLC cell lines. For the murine models, the parental 344SQ-G12D and 393P-G12D cell lines were used, which have the KRAS^G12D^ mutant allele, and using CRISPR-Cas9-HDR gene editing, we generated the KRAS^G12C^ mutant variants, 344SQ-G12C and 393P-G12C cells. For human NSCLC models, H23 (KRAS^G12C^) and A427 (KRAS^G12D^) cells were used. To generate adaptive KRASi-resistant models, the G12C cell lines were treated with MRTX849 (adagrasib, G12Ci) and the G12D lines were treated with MRTX1133 (G12Di). The cells were maintained in incrementally increasing concentrations of the specific KRASi until a stable, surviving, resistant population was generated (Fig. 1A). Growth inhibition assays ascertained the KRASi-resistant cells to have achieved more than 3–10x IC50 values for the specific KRASi as compared to their sensitive parental controls (Fig. 1B and Supp. Fig. S1A). Colony formation assays also confirmed that the KRASi-resistant cells were significantly more resistant to the specific KRAS inhibitors as compared to the sensitive parental cells (Fig. 1C-D). To determine whether the KRASi still blocks the KRAS signaling in the resistant cells, we performed western blot analyses to detect MAPK pathway proteins. We observed that in most of the resistant cells, KRASi treatment resulted in a substantial decrease of pERK and pMEK levels when compared to untreated cells, although the magnitude of suppression was less than in the respective sensitive cells upon KRASi treatment (Fig. 1E and Supp. Fig. S1B). Since we observed that the KRASi-resistant cells still exhibited response to KRAS inhibition, indicating persistent on-target engagement, we sought to understand whether bypass signaling pathways were activated, potentially providing a survival advantage to the resistant cells. For this, we performed proteomic profiling by RPPA (reverse phase protein array) analysis of the resistant/sensitive cells with or without the respective KRASi treatment. We observed that in most of the murine and human KRASi-resistant cells, there was a significant elevation in phospho-PDK1 (3-phosphoinositide-dependent kinase-1) expression upon treatment with the respective KRASi or when compared to the sensitive parental cells (Fig. 1F and Supp. Fig. S1D-E). To validate the finding of the RPPA analyses, western blots were performed, which demonstrated that both in the murine and human cells, the KRASi-resistant cells had elevated expression of total and phosphorylated PDK1 at baseline and upon treatment with KRASi at 24 or 48 hours when compared to the parental sensitive cells (Fig. 1G and Supp. Fig. S1C).

**Figure 1.**
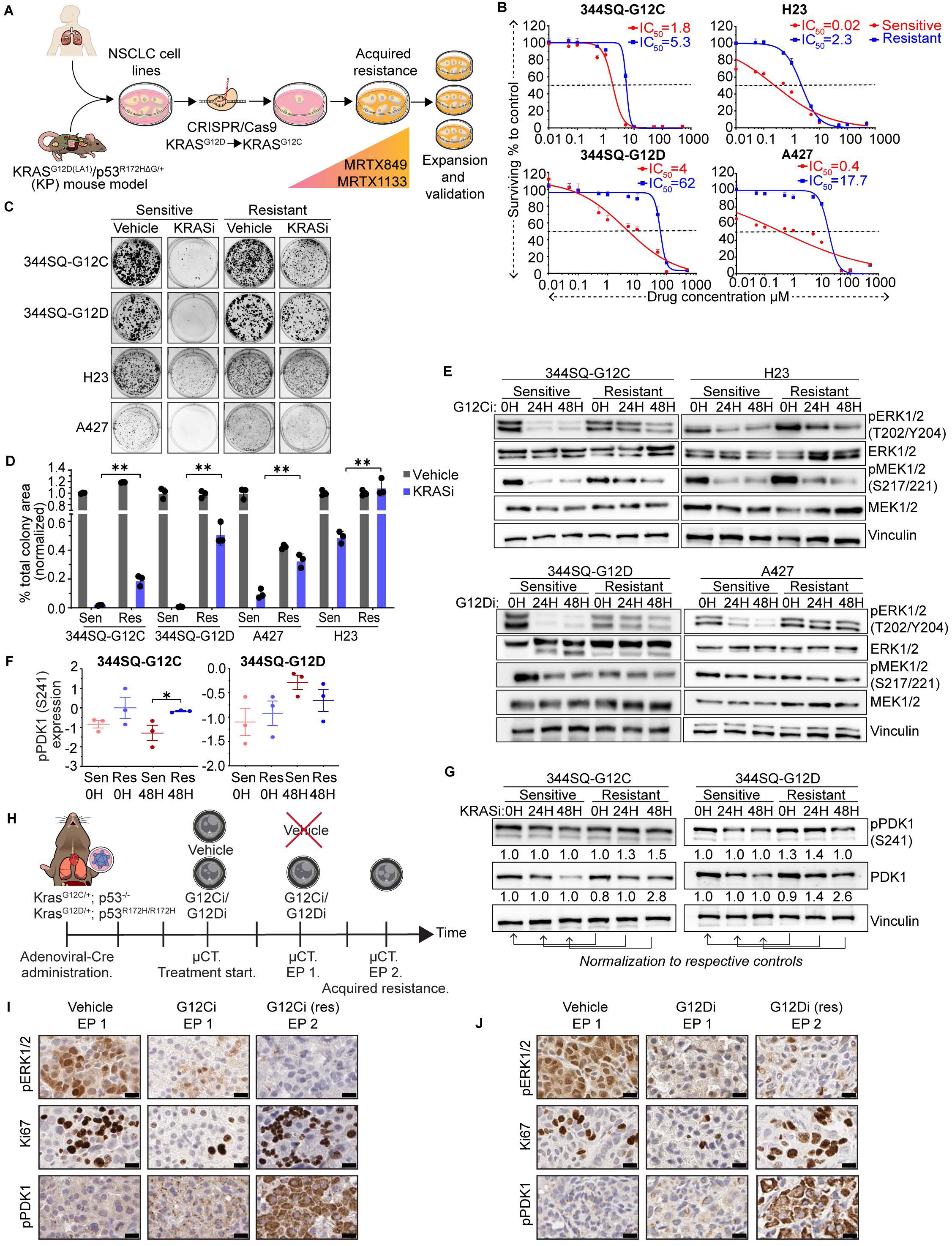
Increased activation of PDK1 in NSCLC with acquired resistance to KRAS inhibition. (A) Schematic demonstrating the workflow of generating KRAS-inhibition (KRASi)-resistant NSCLC cell lines. (B) Growth inhibition assays of KRAS^G12C^ (344SQ-G12C and H23) and KRAS^G12D^ (344SQ-G12D and A427) sensitive (red curve) and resistant (blue curve) cell lines when treated with MRTX849 (G12Ci) and MRTX1133 (G12Di) at the indicated concentrations. (C) Colony formation assay of sensitive and resistant cell lines treated with either DMSO (grey bars) or allele-specific KRAS inhibitors (blue bars) for 72 hours. **, p<0.01. (D) Quantification of the colony formation assay in C. **, p<0.01. (E) Western blotting of sensitive and KRASi resistant lines treated with KRAS inhibitors at IC50 concentrations for the indicated time points to evaluate levels of MAPK pathway components. (F) Dot plots of phosphorylated PDK1 (S241) levels from RPPA analysis of 344SQ-G12C/D sensitive and resistant lines treated with KRAS inhibitors at respective IC50 concentrations for the indicated time points. Statistical significance was calculated using Student’s t-test. *, p<0.05. (G) Western blot analysis of phosphorylated and total PDK1 levels in KRASi sensitive and resistant murine lines treated with MRTX849 or MRTX1133 at IC50 concentrations. (H) Schematic showing generation of genetically engineered mouse (GEM) KRASi resistant models. Adenoviral-Cre was administered to KRAS^LSL-G12C/+^; p53^f/f^ and KRAS^LSL-G12D/+^; ^p53wm/wm^ mice and tumors were tracked using micro-CT imaging. After the tumors became measurable, mice were randomized into vehicle or KRASi treatment groups. The mice were treated until the tumor burden of the vehicle cohorts had reached the humane endpoint, at which time most of the mice were sacrificed (Endpoint, EP 1). A few mice on the KRASi treatment arms continued to be treated until the tumors had regrown (EP 2). (I) IHC staining to evaluate expression of pERK1/2 (T202/Y204), Ki67, and pPDK1 (S241) in tumor nodules from lungs of vehicle and MRTX849-treated mice at EP 1 and 2. Scale bars correspond to 20 µM. (J) IHC staining to evaluate expression of pERK1/2 (T202/Y204), Ki67, and pPDK1 (S241) in tumor nodules from lungs of vehicle and MRTX1133-treated mice at EP 1 and 2. Scale bars correspond to 20 µM.

To establish whether PDK1 signaling was elevated in tumors that acquired KRASi resistance *in vivo*, we used conditional KRAS^G12C/+^;p53^-/-^ or KRAS^G12D/+^;p53^R172H/R172H^ mice^21,22^. Intratracheal delivery of adenoviral-Cre produced autochthonous lung adenocarcinomas with an activated KRAS^G12D^ or KRAS^G12C^ with either homozygous loss of p53 or homozygous expression of p53^R172H^, as indicated by micro-CT imaging (Fig. 1H and Supp. Fig. S1F and S1H). Cohorts of mice were treated with either vehicle as a control or with the respective KRAS inhibitors (MRTX849 for G12C and MRTX1133 for G12D tumors). Periodic micro-CT imaging showed that treatment with KRASi was able to significantly reduce the tumor burden, with maximum response recorded with 57 days (G12Ci) or 42 days (G12Di) of treatment (Endpoint 1) when compared to tumor size at the start of treatment (Day 0). Upon prolonged treatment for 150 days (G12Ci) or 153 days (G12Di) (Endpoint 2), many of the regressed tumor nodules re-emerged as acquired KRASi-resistant tumor outgrowth when quantified by micro-CT (Fig. 1H and Supp. Fig. S1F-I). IHC analysis of the tumors revealed that when compared to vehicle-treated tumors, the KRASi-treated tumors at Endpoint 1 (maximum response) demonstrated significant decreases in pERK and Ki67 with no comparable difference in expression of pPDK1. In the acquired resistant tumors at Endpoint 2, though repression of pERK persisted, there was the emergence of Ki67-positive tumor cells and significantly elevated expression of pPDK1 (Fig. 1I-J). These results support that elevated PDK1 signaling occurs in acquired resistance to KRAS inhibitor *in vivo*.

To determine whether PDK1 signaling is crucial for the generation and maintenance of KRASi resistance, we performed growth inhibition assays using the 344SQ-G12D sensitive or MRTX1133-resistant (M1133Res) cells with combination treatment of a pharmacological PDK1 inhibitor (BX795) and the KRASi. The sensitive cells demonstrated an increased sensitivity to the KRASi when PDK1 was co-inhibited, while the resistant cells showed a dramatic re-sensitization to MRTX1133 when PDK1 was co-targeted, as indicated by the decreased IC50 values (Fig. 2A-B). BX795 alone had no significant differential effect on cell viability of the sensitive or resistant cells (Supp. Fig. S2A). We validated that BX795 treatment functionally inhibited PDK1, as treated cells had reduced levels of pPDK1, as well as pS6 and pRSK2 as downstream target PDK1 activity (Fig. 2C). Colony formation assays also confirmed that the KRASi-resistant cells were significantly re-sensitized to the KRASi upon combination treatment with the PDK1 inhibitor (Fig. 2D-E). To further confirm our observations, we performed similar assays using a second PDK1 inhibitor, GSK2334470, which functionally inhibited PDK1 (Supp. Fig. S2E). Growth inhibition and colony formation assays showed significant combinatorial effects of treatment with the KRASi and GSK2334470 in both the sensitive and resistant 344SQ-G12D cells (Supp. Fig. S2B-D, S2F-G). We observed comparable results with the MRTX849-resistant 344SQ-G12C cells, where the resistant cells were completely re-sensitized to MRTX849 when treated with the combination PDK1 inhibitor (reduced IC50 values in growth inhibition assay and reduced colony formation of M849 resistant cells) (Supp. Fig. S2H-L).

**Figure 2.**
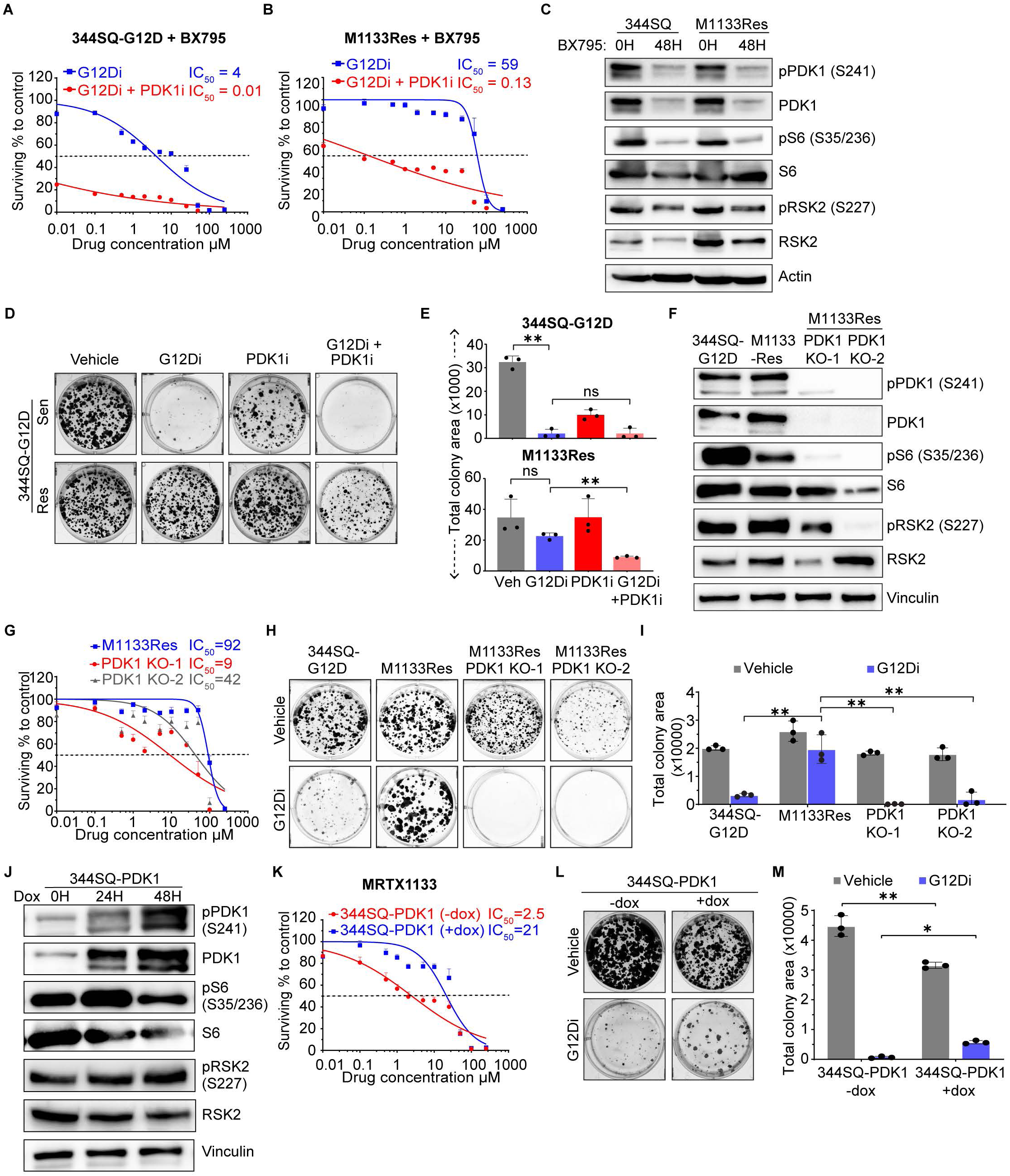
PDK1 expression is necessary and sufficient for generating resistance to KRAS^G12D^ inhibition. **(A)** Growth inhibition assays of 344SQ-G12D cells treated with MRTX1133 (blue line) or co-treated with BX795 at its IC50 concentration (red line). IC50 values for MRTX1133 are indicated on the graph. **(B)** Growth inhibition assays of 344SQ-G12D MRTX1133-resistant line (M1133Res) treated with MRTX1133 (blue line) or co-treated with BX795 at its IC50 concentration (red line). **(C)** Western blot analysis of phosphorylated and total PDK1, S6, and RSK2 in 344SQ-G12D and M1133Res cells upon BX795 treatment at IC50 concentrations (3uM and 8uM, respectively). Actin was used as a loading control. **(D)** Colony formation assays of 344SQ-G12D and M1133Res cells upon treatment with vehicle, MRTX1133, BX795, or both. **(E)** Graph showing quantification of the data in D. *, p<0.05. **, p<0.01. **(F)** Western blot of 344SQ-G12D, M1133Res (parental control), and two PDK1 knockout clones (PDK1 KO-1 and -2) of M1133Res cells to evaluate phosphorylated and total PDK1, S6, and RSK2 levels. **(G)** Growth inhibition assays of M1133Res (blue), PDK1 KO-1 (red) and PDK1 KO-2 (grey) cells upon treatment with MRTX1133. **(H)** Colony formation assays of 344SQ-G12D, M1133Res, and PDK1 KO-1 and KO-2 cells when treated with vehicle or MRTX1133 at the IC30 concentration of the parental control line for 72 hrs. **(I)** Quantification of the colony formation assay in H. **, p<0.01. **(J)** Western blot of 344SQ-G12D cells transduced to stably express PDK1 (344SQ-PDK1) under a doxycycline-inducible promoter. The cells were treated with dox for the indicated time points before collection. The blots were probed for phosphorylated and total PDK1, S6, and RSK2 levels. **(K)** Growth inhibition assays of 344SQ-PDK1 cells with (blue line) or without (red line) 48 hours of dox pretreatment. The IC50 of MRTX1133 for each cell line is indicated on the graph. **(L)** Colony formation assays of 344SQ-PDK1 cells with or without 24 hours of dox treatment. The cells were treated with either DMSO or MRTX1133 at the IC30 concentration of 344SQ-PDK1 +dox cells. The colonies were then allowed to grow out for 3 days in drug-free complete media after which they were stained. **(M)** Quantification of the data in L. *, p<0.05, **, p<0.01.

To further establish the functional necessity of PDK1 signaling in adaptive KRASi resistance, we generated CRISPR/Cas9-mediated genetic PDK1 knockout cell clones using the MRTX1133-resistant M1133Res cells. Western blot analysis confirmed the loss of PDK1 expression in both the clones and reduced levels of downstream pS6 and pRSK2, indicating loss of PDK1 signaling (Fig. 2F). Growth inhibition assays with MRTX1133 revealed a complete re-sensitization of the M1133Res-PDK1 knockout clones (KO-1 and KO-2) as indicated by the reduction of their IC50 values by 2-10-fold (Fig. 2G). Colony formation assay confirmed the significant re-sensitization of the PDK1 KO cells to KRASi as compared to the M1133Res cells (Fig. 2H-I). We next investigated whether PDK1 expression was sufficient to induce resistance to KRASi treatment. For this we generated doxycycline (dox)-inducible PDK1 overexpression in the inherently MRTX1133-sensitive 344SQ-G12D cells. PDK1 expression increased with time of dox induction with subsequent elevation in pS6 levels by 24 hours and elevation in pRSK2 by 48 hours (Fig. 2J). Growth inhibition and colony formation assays performed with MRTX1133 treatment showed that ectopic PDK1 expression in 344SQ-G12D cells produced acquired resistance to MRTX1133, with an 8-fold increase in IC50 values and a significant increase in total colony area (Fig. 2K-M) compared to the control uninduced cells. As PDK1 is canonically downstream of PI3K and could regulate AKT1 ^23–25^, we tested whether inhibiting PI3K or AKT independently could affect acquired resistance to KRASi. Growth inhibition assays performed on the M1133Res cells with combination treatment of KRASi and either a PI3K or AKT inhibitor individually demonstrated little effect to re-sensitize the cells to KRASi as compared to the combination treatment with GSK2334470 (PDK1i) (Supp. Fig. S2M). Additionally, we did not observe any considerable elevation in pAKT levels in the resistant murine or human cells when compared to the sensitive cells upon KRASi treatment (Supp. Fig S2N). These results suggest a potent functional necessity and sufficiency of PDK1 expression in the development of acquired resistance to KRASi that is PI3K/AKT independent.

Next, we wanted to determine if the combination treatment of the KRASi and the PDK1i, *in vivo,* could alter the tumor growth of KRASi-resistant tumors or could increase the magnitude of response of the KRASi. For this, we used our syngeneic mouse models and implanted either the sensitive parental 344SQ-G12D cells or the MRTX1133-resistant (M1133Res) cells in their syngeneic host mice (129/Sv). After initial tumor establishment, animals were divided into equal cohorts and treated with either vehicle, KRASi (MRTX1133) or PDK1i (BX795) alone or with a combination of the inhibitors. Tumor growth curves indicated that in sensitive tumors (344SQ-G12D) the effect of the KRASi alone or the combination showed a similar effective response of significant tumor growth inhibition (TGI). Whereas in the M1133Res tumors, only the combination treatment cohort showed a significant reduction of tumor growth over time, as compared to accelerated tumor growth of the other cohorts (Fig. 3A-B). Accordingly, at the termination of the experiment, the final tumor volumes of the combo group were significantly lower than those of the vehicle or KRASi single-line treatment cohorts (Supp. Fig. S3A). Pharmacodynamic (PD) analysis of the tumors revealed a significant reduction of pERK and Ki67 levels in the KRASi and combo-treated sensitive (344SQ-G12D) tumors. pRSK2 expression was substantially decreased in the combo group, indicating effective inhibition of PDK1 signaling. In the M1133Res tumors though pERK was significantly inhibited both in the KRASi and combination treated tumors, Ki67 was only reduced in the combination group. This indicates that although KRASi alone was able to suppress the MAPK signaling in the resistant tumors, tumor growth was unaffected because of possible dependence on PDK1 signaling. pRSK2 levels were decreased in the single-agent and combo-treated tumors (Fig. 3B). To further validate our observations on the necessity of PDK1 for acquired resistance to KRASi, we utilized the PDK1 knockout cell line that was generated using the MRTX1133-resistant 344SQ-G12D cells. The PDK1-KO-2 cell line was implanted in syngeneic mice along with its parental MRTX1133-resistant 344SQ-G12D cells and the sensitive 344SQ cells. After tumor establishment, each cohort was divided to be treated with either vehicle control or with MRTX1133. The vehicle-treated PDK1-KO cells demonstrated significant TGI and reduction in final tumor volumes as compared to the M1133Res tumors with or without KRASi treatment, with tumor growth and final tumor volumes comparable to the 344SQ sensitive tumors. The KRASi-treated PDK1-KO-2 exhibited significant TGI and reduction of final tumor volumes when compared to their parental M1133Res cells, with tumor growth curves and final tumor volumes matching the 344SQ sensitive cells with KRASi treatment. This indicated that PDK1 loss completely reversed the acquired resistance to the KRASi. This further established that PDK1 expression was also necessary for maintaining KRASi resistance (Fig. 3C-D). Pharmacodynamic (PD) analysis of the tumors revealed a significant reduction of pERK in the KRASi-treated tumors, indicating on-target activity of the inhibitor. Ki67 was only reduced in the KRASi-treated cohorts of the 344SQ-G12D and M1133Res-PDK1-KO2 tumors, whereas the M1133Res tumors did not show any reduction of Ki67 upon KRASi treatment. This indicated that the resistant tumors were able to proliferate by complementary mechanisms, possibly through activation of PDK1 signaling. pRSK2 expression was also completely lost in the PDK1-KO tumors, indicating loss of PDK1 signaling. These results indicate that loss of PDK1 signaling in the M1133Res-PDK1-KO tumors made the M1133-resistant cells revert their resistance to the KRASi, establishing the critical necessity of PDK1 signaling in inducing KRASi resistance (Fig. 3E).

**Figure 3.**
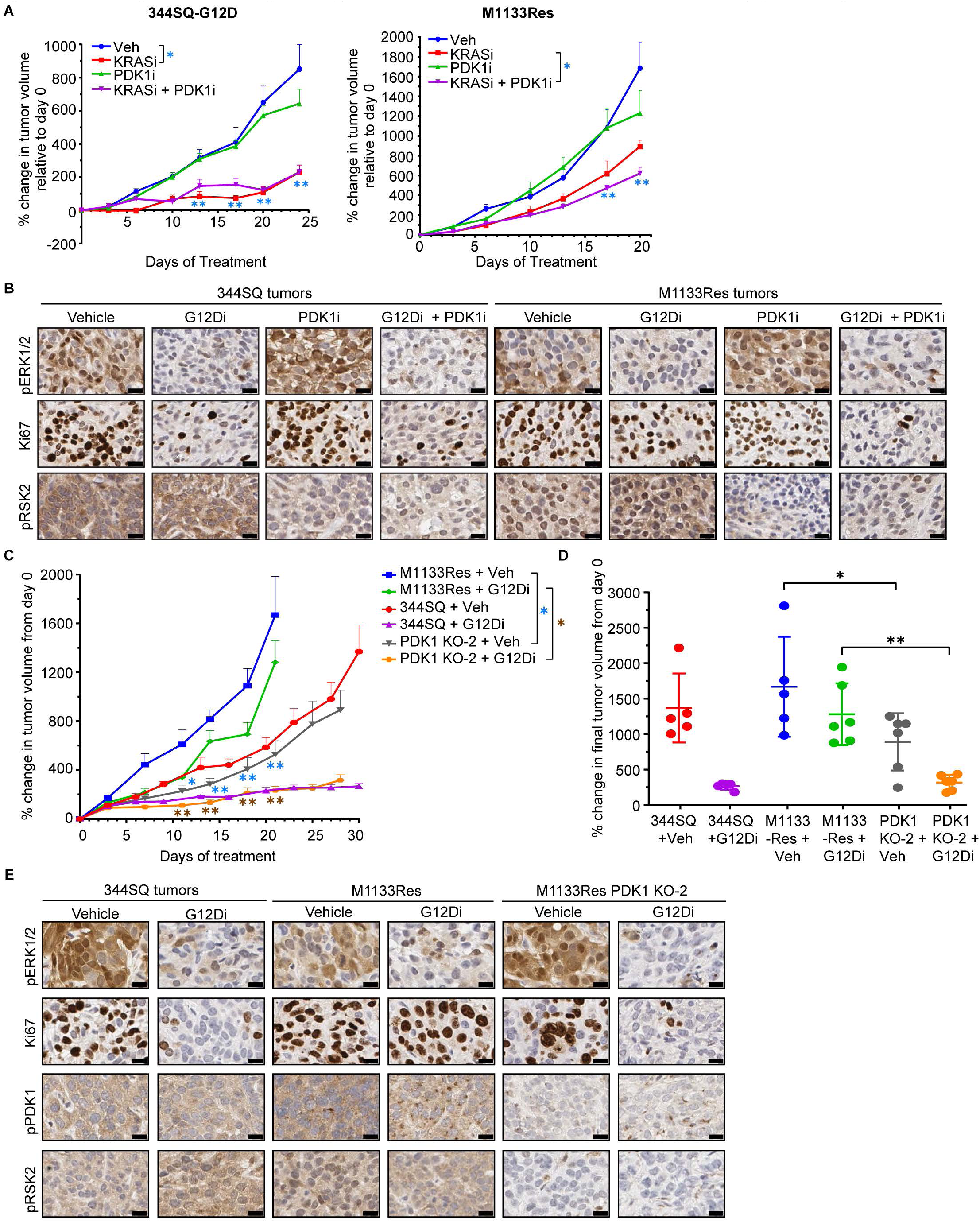
Combination treatment with G12Di and PDK1i results in tumor growth inhibition *in vivo*. **(A)** Mice implanted with 344SQ-G12D (left panel) or M1133Res cells (right panel) were treated with either vehicle (blue line), MRTX1133 (red line), BX795 (green line), or MRTX1133 and BX795 (purple line) for approximately 3 weeks. Statistical comparisons are shown for 344SQ-G12D tumors treated with vehicle vs MRTX1133 (blue asterisks) and M1133Res tumors treated with MRTX1133 vs dual combination (red asterisks) at the indicated time points. **, p<0.01. **(B)** IHC staining is shown for pERK (T202/Y204), Ki67, and pPDK1 (S241) levels in representative sections of tumors harvested from the mice at endpoint. Scale bars correspond to 20 uM. **(C)** Mice implanted with 344SQ-G12D, M1133Res, or M1133Res-PDK1-KO-2 cells. Vehicle-treated tumor growth was recorded for 344SQ (red), M1133Res (blue), and M1133Res-PDK1-KO-2 (grey). MRTX1133-treated tumor growth was recorded for 344SQ (purple), M1133Res (green), and M1133Res-PDK1-KO (orange). Treatment was continued for 4 weeks. Statistical comparisons are shown for M1133Res and M1133Res-PDK1-KO1 cohorts when treated with vehicle (blue asterisks) or MRTX1133 (brown asterisks) at the indicated time points. *, p<0.05. **, p<0.01. **(D)** Percent change in tumor volumes of 344SQ-G12D, M1133Res, and M1133Res-PDK1-KO-2 cohorts from figure 3C at endpoint. Statistical significance was calculated using Student’s t-test. *, p<0.05. **, p<0.01. **(E)** IHC staining is shown for pERK (T202/Y204), Ki67, pPDK1 (S241), and pRSK2 (S227) in representative sections of tumors harvested from mice at endpoint in C. Scale bars correspond to 20uM.

To further validate the *in vivo* efficacy of this KRASi/PDK1i combination, we utilized the conditional KRAS^G12D/+^;p53^R172H/R172H21,22^ GEM model. As described earlier, intratracheal delivery of adenoviral-Cre produced autochthonous lung adenocarcinomas with an activated KRAS^G12D^ and homozygous expression of p53^R172H^, as indicated by Micro-CT imaging. Mice were divided into equal cohorts and treated with either vehicle, MRTX1133 or GSK2334470 (PDK1i) alone as controls or with MRTX1133 + GSK2334470 combo. Imaging after 51 days of treatment showed that KRASi alone or combo treatment was able to significantly reduce tumor burden (Supp. Fig. S3B-C). PD analysis of residual tumors revealed that pERK and Ki67 levels were significantly reduced in the KRASi and combo-treated tumors, with a significant reduction of pPDK1 levels in the combo-treated tumors as compared to the single line treatment cohorts (Supp. Fig.S3D). These results consolidate our in vitro observation and suggest a robust role of combining PDK1i with KRASi for effective TGI of KRASi-resistant tumors *in vivo*.

To elucidate how PDK1 could induce acquired resistance to KRASi, we went back to the RPPA data and searched for activation of known PDK1 downstream pathways. One of the prominent hits we identified was YAP1 activation, which has been reported in different contexts to function downstream of PDK1 ^26–28^. Our data indicated that across murine and human KRASi-resistant models, pYAP1(S127) was elevated when compared with the sensitive cells (Supp. Fig.S4A). Western blot analysis with murine G12C and G12D models validated a consistent elevation of pYAP1 in the resistant cells when compared to the sensitive cells, either at baseline or upon treatment with MRTX849 or MRTX1133 respectively (Fig. 4A). To understand if PDK1 activation was functionally upstream of YAP1 activation in the KRASi-resistant cells, we performed western blot analysis using either the M1133Res-PDK1-KO or the 344SQ-PDK1 cells with dox-induced overexpression of PDK1 and their respective parental or vector control cells. In the M1133Res-PDK1-KO cells, pYAP1 activation was completely abrogated upon KRASi treatment as compared to treated parental cells. Conversely, pYAP1 levels were elevated concurrently with induction of PDK1 expression in the 344SQ-PDK1 cells (Fig. 4B). As the functional effects of YAP1 occur upon nuclear translocation, we further ascertained YAP1 localization in the KRASi-resistant cells by immunofluorescent (IF) staining and quantification, to detect nuclear YAP1 levels in the murine KRASi-resistant and -sensitive cells with or without KRASi treatment. Both the G12C (MRTX849Res) and G12D (M1133Res) resistant murine models demonstrated more cells with elevated nuclear YAP1 as compared to their respective sensitive cells, upon treatment with KRASi (Fig. 4C-D). These results were corroborated in the human KRASi-resistant models, where western blot analysis showed that in comparison to the respective sensitive cells, the KRASi (MRTX849 or MRTX1133) treated H23 or A427 resistant cells had significantly higher levels of the pYAP1-Y357, which is the post-translational modified YAP1 that is specifically associated with nuclear translocated YAP1 (Supp. Fig. S4B). Next, we wanted to validate whether KRASi-resistant cells with higher nuclear YAP1 were functionally activating the TEAD-mediated transcriptional targets. For this, we performed qRT-PCR analysis to detect transcriptional upregulation of a broad list of published TEAD targets. Our results indicated that many of the TEAD target genes were significantly upregulated in the KRASi-resistant cells as compared to the sensitive cells at baseline, as well as in the sensitive cells when treated with the KRASi, when compared to untreated cells (Fig. 4E). These results also indicate a functional activation of YAP1/TEAD in KRASi-resistance. Finally, to determine whether activation of YAP1 is associated with *in vivo* acquired KRASi-resistance, we performed IHC staining of the GEMM lung tumor samples as described in Figure 1G-I and Suppl. Figure S1E-H. We observed that the emergent resistant tumors at Endpoint 2 had more abundant nuclear YAP1 when compared to either the LUAD tumors from mice treated with vehicle or the residual LUAD tumors from the mice that were treated with KRASi at Endpoint 1 at maximum response (Fig. 4F).

**Figure 4.**
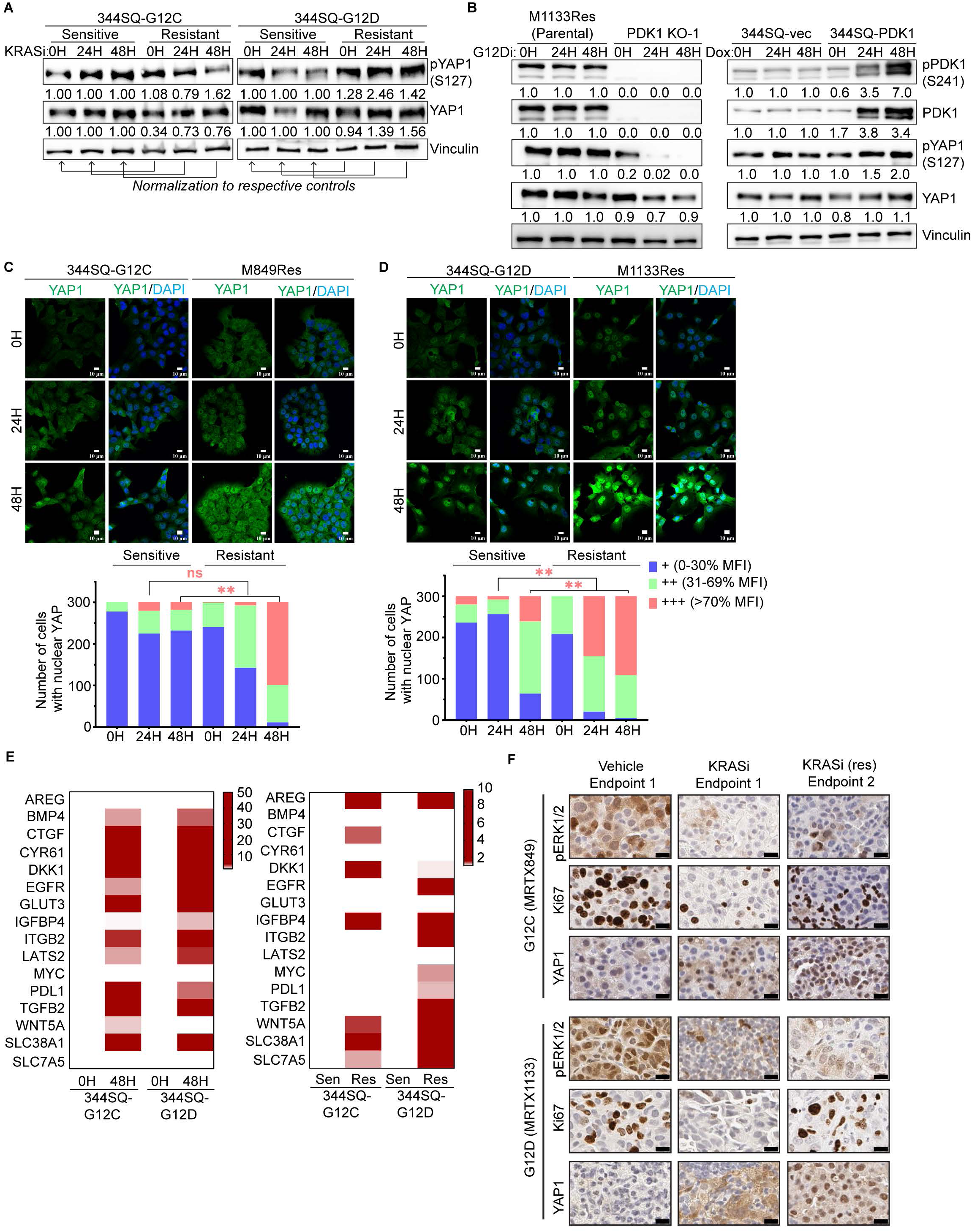
PDK1 functions upstream to activate YAP1/TEAD pathway to drive acquired resistance to KRAS^G12D^ inhibition. (A) Western blot analysis of 344SQ-G12C/D sensitive and KRASi-resistant cells treated with MRTX849 or MRTX1133 at their respective IC50 concentrations. The blots were probed for phosphorylated (S127) and total YAP1 levels. (B) M1133Res (parental control) and PDK1 KO-1 cells were treated with MRTX1133 at IC50 concentrations for the indicated time points and analyzed for levels of phosphorylated and total PDK1 and YAP1 levels using western blotting (left panel). The right panel shows 344SQ-vec (control) and 344SQ-PDK1 lines treated with dox for the indicated time points and probed for expression of phosphorylated and total PDK1 and YAP1. (C) Immunofluorescence staining for YAP1 (green) in 344SQ-G12C and M849Res cells treated with MRTX849 at IC50 concentrations for the indicated time points. Nucleus was stained with DAPI. Scale bars correspond to 10 µM. Quantification of mean fluorescence intensity (MFI) in the nuclei was performed using ImageJ software (lower panel). Triplicates of 100 cells were analyzed for each time point and the number of cells with low (0-30%), medium (31-69%), and high (70-100%) MFI was plotted as indicated. Student’s t-test was used to calculate statistical significance. **, p<0.01. (D) Immunofluorescence staining for YAP1 (green) in 344SQ-G12D and M1133Res cells treated with MRTX1133 at IC50 concentrations for the indicated time points. Nucleus was stained with DAPI. Scale bars correspond to 10 µM. Student’s t-test was used to calculate statistical significance. **, p<0.01. (E) The heatmap on the left shows qPCR analysis to evaluate expression of TEAD transcriptional targets in 344SQ-G12C and 344SQ-G12D cells with or without KRASi treatment. qPCR analysis of 344SQ-G12C and 344SQ-G12D sensitive and resistant lines is shown on the right. Heatmaps were generated using GraphPad Prism. (F) IHC staining to evaluate expression of pERK1/2 (T202/Y204), Ki67, and YAP1 in lung tumor nodules from KRAS^LSL-G12C/+^; p53^f/f^ and KRAS^LSL-G12D/+^; ^p53wm/wm^ mice with acquired resistance to KRASi (figure 1 and supplemental figure 1). Scale bars represent 20 µM.

We conducted growth inhibition assays utilizing 344SQ-G12D sensitive or MRTX1133-resistant cells treated in combination with a pharmacological TEAD inhibitor (VT107) and the KRASi to ascertain if YAP1/TEAD signaling is essential for the development and maintenance of KRASi resistance. When TEAD was co-inhibited, the sensitive cells showed greater sensitivity to the KRASi, while the resistant cells showed substantial re-sensitization to MRTX1133, as seen by lower IC50 values (Fig. 5A-B). VT107 alone had no significant influence on the viability of sensitive cells compared to resistant cells (Supp. Fig. S5A). We confirmed that VT107 treatment functionally suppressed YAP1/TEAD, as treated cells showed lower levels of pYAP1, YAP1 and CYR61 which is a downstream target of TEAD (Fig. 5C). Colony formation assays further revealed that KRASi-resistant cells were greatly re-sensitized to KRASi when combined with the TEAD inhibitor (Fig. 5D–E). To determine whether MRTX849-resistant cells have similar dependency on YAP1/TEAD activation, we performed similar experiments and observed comparable results with MRTX849-resistant 344SQ-G12C cells, which were entirely re-sensitized to MRTX849 when treated with the combination TEAD inhibitor treatment (lower IC50 values in growth inhibition assays and reduced colony formation) (Supp. Fig. S5B-F). To demonstrate the functional importance of YAP1/TEAD signaling in adaptive KRASi resistance, we created CRISPR/Cas9-mediated genetic knockout cell clones for YAP1 utilizing MRTX1133-resistant 344SQ-G12D cells (M1133Res). Western blot analysis verified the loss of YAP1 expression in both clones (Fig. 5F). The growth inhibition assay with MRTX1133 showed that the M1133Res-YAP1 knockout clones (KO-1 and KO-2) were completely re-sensitized to KRASi, as evidenced by a 20-25-fold decrease in their IC50 values (Fig. 5G). The M1133Res-YAP1 KO cells were also significantly more sensitive than the M1133Res cells, according to the results from the colony formation experiment (Fig. 5H-I).

**Figure 5.**
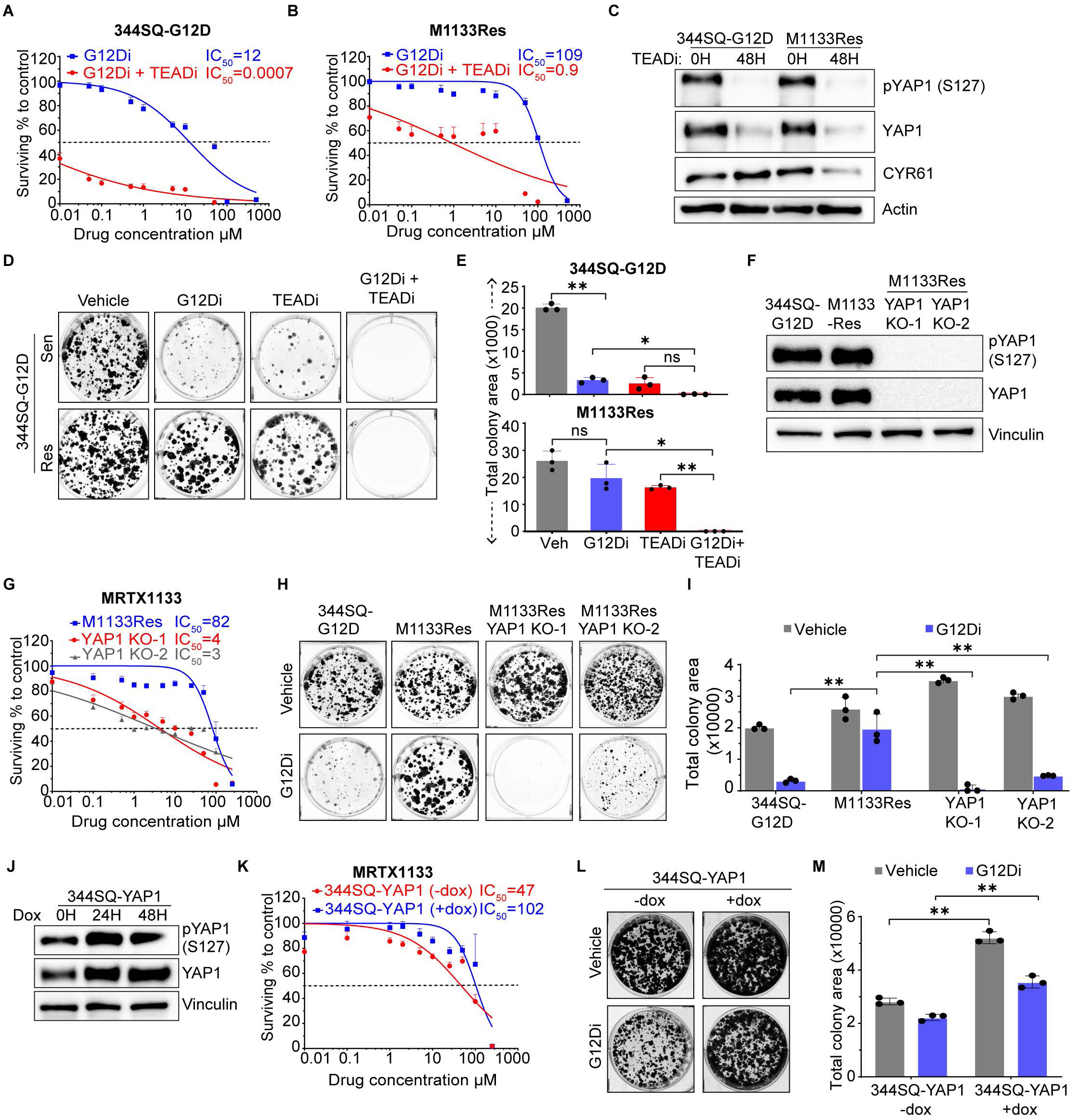
YAP1/TEAD pathway is necessary and sufficient for driving resistance to KRAS^G12D^ inhibition. (A) Growth inhibition assays of 344SQ-G12D cells when treated with serial dilutions of MRTX1133 as indicated (blue line) or co-treated with VT107 (TEADi) at its IC50 concentration (red line). (B) Growth inhibition assays of M1133Res cells treated with MRTX1133 at indicated concentrations (blue line) or co-treated with VT107 at its IC50 concentration (red line). (C) Western blot of 344SQ-G12D and M1133Res cells when treated with VT107 at IC50 concentrations for the indicated time points to examine pYAP1 (S127), YAP1, and CYR61 levels upon TEAD inhibition. (D) Colony formation assays of 344SQ-G12D and M1133Res cells treated with the indicated drugs for 72 hrs. (E) Quantification of the data in D. *, p<0.05. **, p<0.01. (F) Western blotting to confirm two YAP1 knockout clones generated from M1133Res cells as evidenced by pYAP1 (S127) and YAP1. (G) Growth inhibition assays of M1133Res (blue), YAP1 KO-1 (red), and YAP1 KO-2 (grey) upon treatment with MRTX1133. (H) Colony formation assays of 344SQ-G12D, M1133Res (control), and YAP1 KO-1 and KO-2 cells after treatment with DMSO or MRTX1133 at IC30 concentration of the control cell line for 72 hrs. (I) Quantification of the data in H. (J) Western blot of 344SQ-G12D cells transduced to stably express YAP1 (344SQ-YAP1) under a dox-inducible promoter. The cells were treated with dox for the indicated time points before lysis. The samples were probed for phosphorylated and total YAP1 levels. (K) Growth inhibition assays of 344SQ-YAP1 cells with (blue) or without (red) 72 hours of dox pretreatment. The cells were treated with MRTX1133 at the indicated concentrations for 72 hrs. (L) Colony formation assays of 344SQ-YAP1 cells following 72 hours of dox treatment. The cells were treated with either DMSO or MRTX1133 for 72 hours at the IC30 concentration of 344SQ-YAP1 +dox cells. The colonies were then allowed to grow out for 3 days in drug-free media after which they were stained. (M) Quantification of the colony formation assay in L. **, p<0.01.

We then sought to determine if YAP1 expression was adequate to induce acquired resistance to KRASi. For this, we produced doxycycline (dox) inducible YAP1 overexpressing 344SQ-G12D cells, which are MRTX1133-sensitive by nature. With increasing duration of dox induction, YAP1 expression also increased (Fig. 5J). Ectopic induction of YAP1 expression in 344SQ-G12D cells produced acquired resistance to MRTX1133, as demonstrated by a 2.2-fold increase in IC50 values in a growth inhibition assay and a significant increase in total colony area (Fig. 5K-M) when compared to the uninduced control cells in colony formation assays conducted with MRTX1133 treatment.

Further, we sought to ascertain whether treating KRASi and TEADi together *in vivo* may change the growth of KRASi-resistant tumors or boost the KRASi’s response. The sensitive parental 344SQ-G12D cells or the MRTX1133-resistant (M1133Res) cells were implanted in their syngeneic host mice (129/Sv). After the initial tumor development, animals were split into equal cohorts and given either vehicle, KRASi (MRTX1133), or TEADi (VT107) alone or a combination of MRTX1133 and VT107. Tumor growth curves revealed that in sensitive tumors (344SQ-G12D), the effect of KRASi alone or the combination of inhibitors produced a comparable effective response of significant tumor growth inhibition. In the M1133Res tumors, only the combination treatment group demonstrated a significant reduction in tumor growth over time, as opposed to the accelerated tumor development of the other cohorts (Fig. 6A-B). As a result, at the end of the experiment, the combined group had considerably lower final tumor volumes than the vehicle or KRASi single-line therapy cohorts (Supp. Fig. S6A). The pharmacodynamic (PD) study of the tumors demonstrated a considerable reduction in pERK and Ki67 levels in KRASi and combo-treated sensitive tumors. In the M1133Res tumors, pERK was significantly reduced in the KRASi alone and the combo treatment cohorts, but Ki67 was only decreased in the combination-treated tumors. Additionally, CYR61 is a transcriptional TEAD target that was robustly suppressed in the tumors treated with either VT107 or the combination. These results indicate a potential role of YAP1/TEAD signaling in the maintenance of KRASi resistance (Fig. 6B). To further establish the *in vivo* efficacy of this combination, we utilized the conditional KRAS^G12D/+^;p53^R172H/R172H^ ^21,22^ GEMM. As described earlier, intratracheal delivery of adenoviral-Cre produced autochthonous lung adenocarcinomas with an activated KRAS^G12D^ and homozygous expression of p53^R172H^, as indicated by Micro-CT imaging. Mice were divided into equal cohorts and treated with either vehicle, MRTX1133 or VT107 (TEADi) alone as controls or with MRTX1133 + VT107 combo. Imaging after 42 days of treatment revealed that KRASi alone or the combo treatment was able to significantly reduce tumor burden (Supp. Fig. S6B-C). PD analysis of residual tumors revealed that pERK and Ki67 levels were significantly reduced in the KRASi and combo-treated tumors, with a significant reduction of CYR61 levels in the VT107 and combo-treated tumors as compared to the vehicle-treated tumors (Supp. Fig.S6D). These results consolidate our in vitro observation and suggest a robust role of combining TEADi with KRASi for effective tumor growth inhibition of KRASi-resistant tumors *in vivo*.

**Figure 6.**
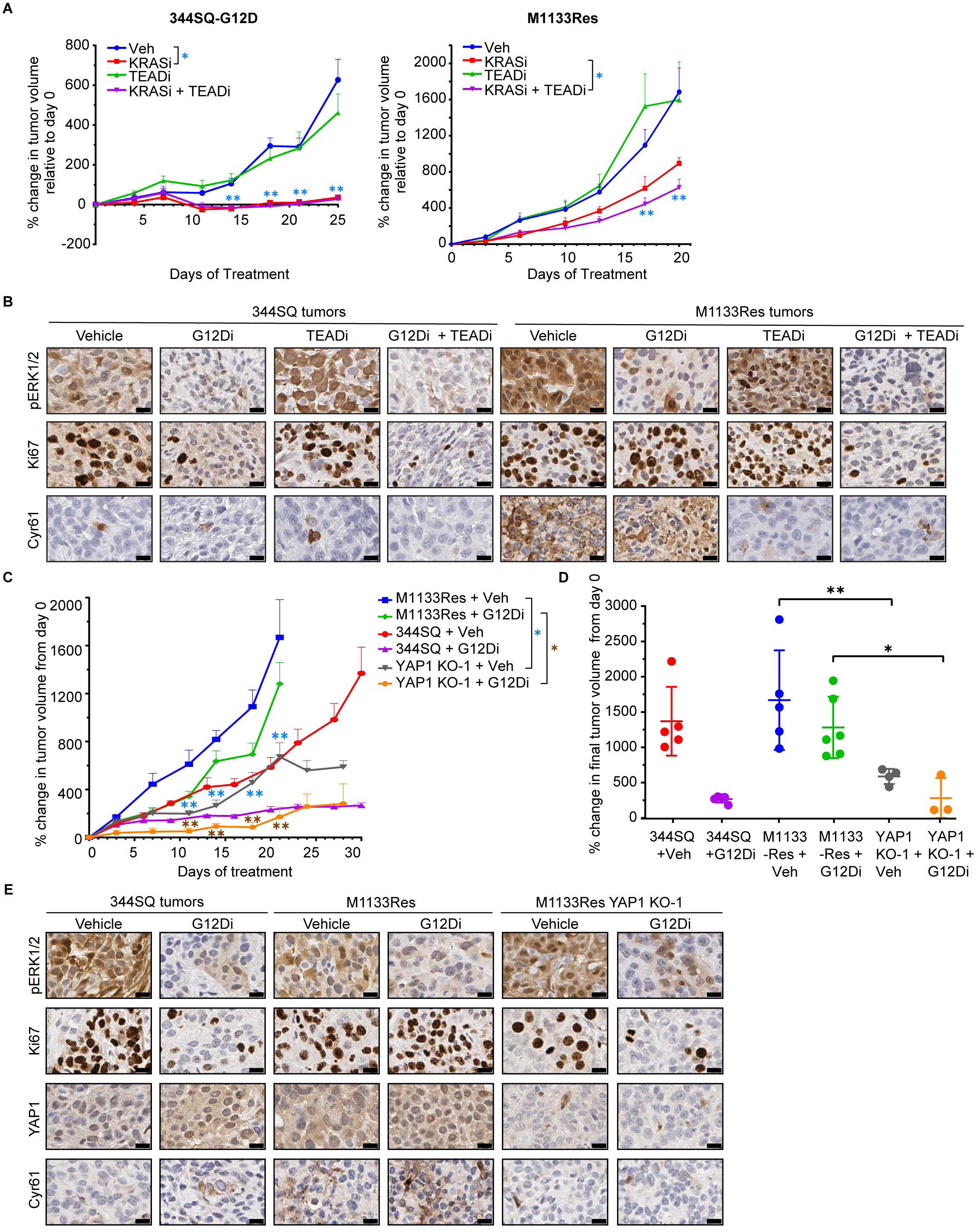
Combination treatment with G12Di and TEADi results in tumor growth inhibition *in vivo*. **(A)** Mice implanted with 344SQ-G12D (left panel) or M1133Res cells (right panel) were treated with either vehicle (blue line), MRTX1133 (red line), VT107 (green line), or MRTX1133 and VT107 (purple line) for approximately 3 weeks. The M1133Res vehicle and KRASi treated cohorts shown here are shared with the experiment in 3A. Statistical significance is shown for 344SQ-G12D tumors treated with vehicle vs MRTX1133 (blue asterisks) and M1133Res tumors treated with MRTX1133 vs combination of MRTX1133 and VT107 (red asterisks) at the indicated time points. Statistical significance was calculated using Student’s t-test. **, p<0.01. **(B)** IHC analysis to evaluate the expression of pERK (T202/Y204), Ki67, YAP1, and CYR61 in subcutaneous tumor sections collected from mice at the end of treatment. Scale bars corresponding to 50 µM are shown. **(C)** Mice implanted with 344SQ-G12D, M1133Res, or M1133Res-YAP1-KO1 cells. Vehicle-treated tumor growth was recorded for 344SQ (red), M1133Res (blue), and M1133Res-YAP1-KO1 (grey). MRTX1133-treated tumor growth was recorded for 344SQ (purple), M1133Res (green), and M1133Res-YAP1-KO1 (orange). Treatment was continued for 4 weeks. Statistical comparisons are shown for M1133Res and M1133Res-YAP1-KO1 cohorts when treated with vehicle (blue asterisks) or MRTX1133 (brown asterisks) at the indicated time points. **, p<0.01. **(D)** Percent change in tumor volumes of 344SQ-G12D, M1133Res, and M1133Res-YAP1-KO-1 cohorts from figure 6C at endpoint. Statistical significance was calculated using Student’s t-test. *, p<0.05. **, p<0.01. **(E)** IHC staining is shown for pERK (T202/Y204), Ki67, YAP1, and Cyr61 in representative sections of tumors harvested from mice at endpoint in C. Scale bars correspond to 20uM.

To further validate our observations on the necessity of YAP1/TEAD signaling for acquired resistance to KRASi, we utilized the YAP1 knockout cell line that was generated using the MRTX1133-resistant 344SQ-G12D cells (M1133Res). The YAP1-KO-1 cell line was implanted in syngeneic mice along with its parental MRTX1133-resistant 344SQ-G12D cells and compared to the sensitive 344SQ parental tumors. After tumor establishment, each cohort was divided to be treated with either vehicle control or with MRTX1133. The KRASi-treated YAP1-KO-1 exhibited significant TGI and reduced final tumor volumes when compared to their parental M1133Res cells, and the 344SQ sensitive cells when treated with KRASi. This indicated that YAP1 loss completely abrogated acquired resistance to the KRASi. The vehicle-treated YAP1-KO cells also demonstrated significant TGI and reduced final tumor volumes as compared to the M1133Res tumors with or without KRASi treatment, with tumor growth and final tumor volumes comparable to the 344SQ sensitive tumors with KRASi treatment. This further established that YAP1 expression was also necessary for maintaining KRASi resistance (Fig. 6C-D). Pharmacodynamic (PD) analysis of the tumors revealed a significant reduction of pERK in the KRASi-treated tumors, indicating on-target activity of the inhibitor. Ki67 was only reduced in the KRASi-treated cohorts of the 344SQ-G12D and M1133Res-YAP1-KO-1 tumors, whereas the M1133Res tumors did not show any reduction of Ki67 upon KRASi treatment. This indicated that the resistant tumors were able to proliferate by complementary mechanisms, possibly through activation of YAP1 and PDK1 signaling. Total YAP1 and Cyr61 expression also completely lost in the YAP1-KO tumors, indicating loss of YAP1/TEAD signaling in the KO tumors, whereas elevated Cyr61 in the M1133Res tumors as compared to the 344SQ-G12D sensitive tumors indicated elevated YAP1/TEAD signaling in the resistant tumors (Fig. 6E). These results indicate that loss of YAP1 in the M1133Res-YAP1-KO tumors made the M1133-resistant cells revert their resistance to the KRASi, establishing the critical necessity of YAP1/TEAD signaling in inducing KRASi resistance.

## Discussion

The development of direct KRAS inhibitors has been a major advancement in the treatment of KRAS-driven epithelial tumors, including NSCLC^29–31^. Like other targeted therapies, prolonged KRASi treatment also results in the emergence of acquired and adaptive resistance mechanisms, which have been observed clinically (for adagrasib and sotorasib)^32,33^ and in preclinical settings for both KRAS mutant allele-specific inhibitors (e.g. MRTX1133, LY3962673, Olomorasib), restraining the durability and efficacy of these novel therapeutic agents^32,34^. Understanding the causation of acquired resistance to KRASi is therefore an urgent necessity for the development of combinatorial therapeutic approaches to either overcome or delay resistance to KRASi. Currently, for NSCLC, it has been demonstrated that resistance to KRAS^G12C^ inhibitors is a result of genetic mutations or amplification of either RAS genes^14,32^ or genes like BRAF and MAP2K1, resulting in reactivation of the MAPK pathway^14,32,34,35^. Amplification of RTKs like EGFR and MET has also been implicated^34,36^. Here, we have used genetically engineered mouse models, established syngeneic cell line models, and human NSCLC cell lines to generate acquired resistance to KRAS^G12C^ and KRAS^G12D^ inhibitors, followed by proteomic profiling to specifically identify activation of complementary signaling pathways that could bypass MAPK activation to impart resistance. Among several other signaling genes that were modulated, we identify PDK1 activation as a novel path to KRASi resistance. Using pharmacological inhibition, genetic gain-of-function and loss-of-function approaches, we demonstrate the critical role of PDK1 activation in driving and maintaining KRAS allele-specific inhibitor resistance. Though PDK1 pathway activation has been implicated in driving resistance to osimertinib in EGFR mutant lung cancers^27^, our report is the first to functionally associate activation of PDK1 with KRASi resistance. Further, our data substantiates the strong functional role of YAP1/TEAD activation in both KRAS^G12C^ and KRAS^G12D^ inhibitor resistance. This finding of YAP1/TEAD activation corroborates previous observations of its role in KRAS ^G12C^ inhibitor resistance^15,20,26,37^. Overall, from our *vitro* and *in vivo* functional assays, we therefore validate the critical necessity and sufficiency of both PDK1 and YAP1/TEAD pathway activation in KRAS inhibitor resistance. Interestingly, we also observe that PDK1 could regulate YAP1/TEAD activation as a downstream effect, as YAP1 activation was completely abrogated in PDK1 knockout M1133Res tumor cells. Also, ectopic expression of PDK1 directly induced YAP1/TEAD pathway activation. This is the first time a direct upstream regulatory pathway for YAP1/TEAD activation in KRASi resistance has been demonstrated, which was unclear in previous literature that elucidated the activation of YAP1/TEAD in KRAS^G12C^ resistance. Currently, we are trying to understand the molecular mechanisms of this regulation of YAP1 by PDK1, which is outside the scope of this manuscript. Our current focus is to understand the functional role of these pathways in regulating the tumor microenvironment. Overall, this paper provides solid pre-clinical evidence for a critical role of PDK1 and the YAP1/TEAD signaling pathway activation in the induction of acquired resistance to KRAS allele-specific inhibitors.

## Materials and Methods

### Cell culture

344SQ and 393P cell lines were generated as described previously ^17^. All other cell lines were purchased from ATCC. HEK293T cells were cultured in DMEM high glucose media (Gibco) supplemented with 10% fetal bovine serum (FBS). All other cell lines were maintained in RPMI 1640 (Gibco). To generate KRAS mutant allelic versions, mouse Kras^G12D^;p53^R172H^ NSCLC cell lines were converted to Kras^G12C^ by CRISPR homologous-direct repair. Approximately 50,000 cells 50,000 cells were plated in a 6-well plate, and endogenous KrasG12D was knocked out using the snRNP complex by co-transfecting 13.8 µg of Cas9 nuclease (IDT) with 2.8 µg of G12D sgRNA (sequence below) and 1 µM of G12C ssODN (sequence below) using Lipofectamine CRISPRMax reagent (Thermo). After 48 hours following transfection, cells were cultured in 1 µM of KrasG12D inhibitor MRTX1133 to select for positively converted cells. sgG12D: 5’-GTGGTTGGAGCTGATGGCGT-3’.

G12C ssODN Sequence: 5’-AGTTGTATTTTATTATTTTTATTGTAAGGCCTGCTGAAAATGACTGAGT ATAA GCTTGTGGTGGTTGGAGCTTGTGGTGTAGGCAAGAGCGCCTTGACGATACAGCTAATTCAGAATCACTTGTGGATGAGTATGACCC-3’.

To generate KRASi-resistant lines, KRASi-sensitive cells were passaged with mutant-specific KRAS inhibitor treatment starting at IC50 concentrations under conditions of sequential dose escalation until the cells were able to survive in at least fivefold higher drug concentrations. To generate PDK1 and YAP1 knockout lines, cells were transfected with Gene Knockout Kits purchased from Synthego (EditCo) targeting the gene of interest using Lipofectamine CRISPRMAX (Invitrogen) according to the manufacturer’s instructions. After transfection, the cells were seeded in very low density until the growth of colonies was observed. The individual colonies were then picked from the plate separately passaged and screened for the loss of target protein using western blotting. To generate PDK1 and YAP1 overexpression lines, full length mouse cDNA (GeneScript) was cloned into the doxycycline-inducible pTRIPZ-GFP vector using methods described previously ^38^. Lentivirus was generated by transfecting lentiviral constructs with packaging plasmids psPAX2 and pMD2.G in HEK293T cells using Lipofectamine 2000 (Thermo Fisher). Cells were infected using 8 µg/mL polybrene and cultured under puromycin selection ^39^. Treatment with a final concentration of 2 µg/mL dox for indicated time periods was performed before the cells were used for experiments. All cell lines were routinely tested for mycoplasma contamination (LookOut Mycoplasma Elimination Kit, Sigma-Aldrich).

### Growth inhibition assays

2500 cells were seeded in each well of a 96-well flat-bottom clear plate and allowed to adhere overnight in complete media. The following day, fresh media was added containing inhibitors at serially diluted concentrations as is indicated in the figures, and the cells were incubated with the drugs for 72 hours. Eight wells were seeded per concentration. DMSO was added to the media for blank wells. Post treatment, the cells were incubated with a tetrazolium-based dye (G4100, Promega) for 4 hours at 37°C in a humidified CO2 chamber. Reaction was stopped by the addition of a solubilization/stop solution which dissolved the formazan product generated by the living cells. Absorbance was recorded at wavelengths of 570 nm and 700 nm (reference wavelength) using BioTek Epoch Microplate reader (Agilent), and the delta of the two values was used to calculate cell viability. The data was plotted as the percent difference in cell viability at each concentration compared to blank. GraphPad Prism was used to calculate the IC50 values.

For single agent growth inhibition assays, the viability of the cells was measured in serially diluted amounts of that agent. For the dual combination viability assays, the IC50 of the combination agent was first worked out for each cell line. The KRAS inhibitor was then serially diluted in media containing the combination agent dissolved at its IC50 concentration for that cell line.

### Colony formation assays

2000 cells per well were plated onto 6-well plates and allowed to adhere overnight. The following day, drugs were added to the growth media at IC30 concentrations, and the cells were allowed to incubate with the drugs for 72 hours. Each condition was run in triplicates. Post incubation, the cells were allowed to grow in fresh media for 3-5 days and the resulting colonies were fixed and stained with crystal violet. Plates were scanned and colony area was measured using ImageJ. Data in each of the graphs is representative of the mean ± SD of n=3 samples. Statistical significance was calculated using two-way ANOVA.

### Western blot

Cells were washed with PBS and lysed using 1x RIPA buffer supplemented with protease (5871, CST) and phosphatase (P5726 and P0044, Sigma-Aldrich) inhibitor cocktails. Samples were sonicated and centrifuged to collect the supernatant. The samples were boiled in 1x Laemmli buffer at 95°C for 5 minutes before being run on polyacrylamide gels. Gels were transferred onto nitrocellulose membranes. All membranes were blocked using 5% nonfat dry milk in 1x TBST. Primary antibody dilutions were made in 3% BSA and 0.05% NaN3 in 1x TBST. Bands were visualized using Bio-Rad ChemiDoc Imagers and quantified using the Bio-Rad Image Lab software. Wherever quantification is shown, the band intensities were normalized to their respective loading control. Primary antibodies are listed in supplemental table 1.

### RPPA

Cells were washed with PBS and lysed using a lysis buffer (1% Triton X-100, 50 mM HEPES [pH 7.4], 150 mM NaCl, 1.5 mM MgCl2, 1 mM EGTA, 100 mM NaF, 10 mM NaPPi, 1 mM Na3VO4, 10% glycerol and freshly added protease and phosphatase inhibitors from Roche [#05056489001 and #04906837001] made in ddH2O. The samples were then incubated on ice for 20 minutes and centrifuged at 4°C to collect the supernatant. Each sample was submitted to the RPPA core in triplicates. The extracted protein was boiled in 1x SDS sample buffer and submitted for RPPA as described previously ^40,41^.

### Immunohistochemistry

All immunohistochemical staining was performed on formalin-fixed paraffin-embedded tissue sections. Heat-mediated antigen retrieval was performed using either a sodium citrate or a Tris-EDTA buffer, and sections were blocked using 5% goat serum. Overnight incubation at 4°C was performed with primary antibodies at dilutions ranging from 1:200-1:500. Sections were incubated with horseradish peroxidase conjugated secondary antibody (SignalStain, CST) for one hour at room temperature (RT) before staining with DAB. All primary antibodies used are listed with their dilutions in supplemental table 1. Slides were scanned using Aperio ScanScope Turbo slide scanner (Leica Biosystems) and viewed using ImageScope software.

### RNA isolation and qRT-PCR

RNA extraction was performed using Trizol reagent (Invitrogen) according to manufacturer’s instructions. RNA was used for cDNA synthesis or stored in -80. cDNA was isolated using qScript cDNA synthesis kit (Quantabio), followed by qPCR with iTaq Universal SYBR Green Supermix (Bio-Rad). L32 expression was used as the housekeeping gene. The qPCR reactions were performed on the Bio-Rad CFX384 Real-Time System. Primers used are listed in supplemental table 2.

### Cell immunofluorescence

Cells grown on glass cover slips were washed with PBS and fixed with 4% paraformaldehyde (PFA) at room temperature. Cells were permeabilized with 0.1% Triton X-100 and blocked with 5% goat serum. Primary antibody incubation was performed overnight at 4°C followed by fluorophore-tagged secondary antibody (A11008, Invitrogen) incubation for one hour at RT. Counterstaining was performed with DAPI and images were acquired on a confocal microscope. Mean fluorescence intensity (MFI) was quantified using ImageJ.

### Animal experiments

All animal studies were performed with the approval of the University of Texas MD Anderson Cancer Center Institutional Animal Care and Use Committee (IACUC). In the syngeneic xenograft experiments, cells were resuspended in 100 µL serum-free RPMI and subcutaneously implanted under the right flank of 129/Sv male mice of age 3 to 6 months. For the experiments in figures 3A and 6A, 1.5x10^6^ cells were injected in the 344SQ-G12D cohorts, and 0.75x10^6^ cells were implanted in the M1133Res cohorts. Tumors were measured twice a week using calipers, and treatment regimens were initiated once tumors had reached volumes of 50-75 mm^3^. Statistical comparisons for all the xenograft experiments were performed using Student’s t-test.

For the GEMM experiments, mice were intubated with adenovirus Cre at a titer of 3 x 10^7^ pfu. Micro-CT imaging was performed monthly to monitor tumor formation. After confirming presence of tumors, mice were randomized into cohorts and treatment regimens initiated. To analyze tumor growth, three tumor nodules were tracked per mouse using micro-CT images and tumor area was quantified using ImageJ.

### Drug treatments

MRTX849 and MRTX1133 (Mirati) were dissolved in 10% Captisol at 10 mg/mL and 3 mg/mL, respectively. MRTX849 was administered orally at 100 mg/kg QD, and MRTX1133 was injected via i.p. injections at 30 mg/kg ^7,42^. BX795 (Selleck Chemicals), VT107 and GSK2334470 (MedChemExpress) were dissolved in a PEG-based solvent according to the manufacturer’s instructions. For the combination experiments with MRTX1133 and the PDK1 inhibitors, mice were given each drug on alternate days for the duration of the treatment such that each mouse received one dose of a drug per day. BX795 was given at 20 mg/kg and GSK2334470 was given at 80 mg/kg intraperitoneally ^43,44^. For the combination experiments with MRTX1133 and VT107, MRTX1133 was given daily and 10 mg/kg VT107 was administered orally on alternate days. Each dose was given at a volume of 300 uL.

## Author’s Contributions

Study conceptualization, design, and implementation of project: A.B, S.T.K, and D.L.G. Development of cell lines: A.B., S.T.K, B.S.C, and D.H.P. Data acquisition and analysis: A.B, S.T.K, B.S.C, S.G.G, and E.M. Statistical analysis, interpretation, and representation of data: A.B, S.T.K, B.S.C, L.D, and J.W. Manuscript writing, critical revisions, and preparation of figures and tables: A.B, S.T.K, J.J.F, and D.L.G. Overall supervision and execution: S.T.K and D.L.G.

### Acknowledgements

Data was generated in part through the use of the Research Animal Support Facility (RASF) and the Functional Proteomics Reverse Phase Protein Array Core, which receives partial support from the National Cancer Institute under grant P30CA016672 to MD Anderson Cancer Center and Dr. Yiling Lu’s NIH R50 Grant #R50CA221675. The research reported in this paper was not directly funded through the grant P30CA016672 to MD Anderson Cancer Center and is not within the scope of such a grant. We thank Eileen McCarthy for technical assistance with some animal experiments.

---

**Supplemental Figure S1.**
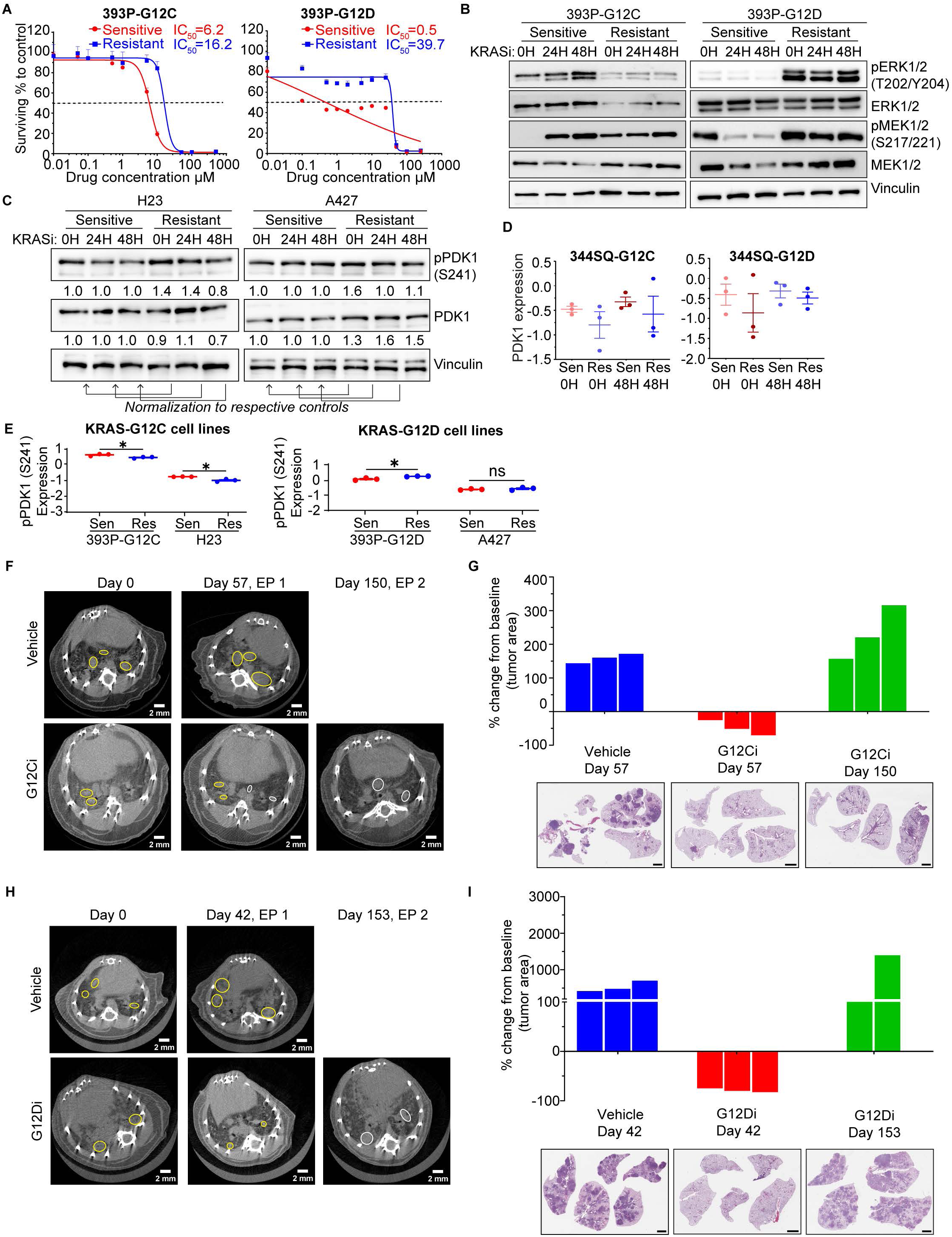
Increased PDK1 activation in KRASG12C-driven NSCLC with acquired resistance to MRTX849. (A) Growth inhibition assays of 393P-G12C (left) and -G12D (right) sensitive and resistant cell lines under treatment with MRTX849 and MRTX1133. (B) Western blotting of 393P-G12C/D sensitive and resistant lines treated with MRTX849 and MRTX1133, respectively, at IC50 concentrations for the indicated time points. The blots were analyzed for levels of phosphorylated and total ERK1/2 (T202/Y204) and MEK1/2 (S217/221). (C) Western blot analysis of H23 and A427 sensitive and resistant lines when treated with KRASi at IC50 concentrations for the indicated time points to evaluate pPDK1 (S241) and PDK1 levels. (D) Dot plots of total PDK1 levels from RPPA analysis of 344SQ-G12C/D sensitive and resistant lines treated with KRAS inhibitors at respective IC50 concentrations for the indicated time points. (E) Dot plots of PDK1 phosphorylation (S241) from RPPA analysis of two KRAS^G12C^ and KRAS^G12D^ sensitive and resistant lines treated with allele-specific KRAS inhibitors at respective IC50 concentrations for the indicated time points. Statistical significance was calculated using Student’s t-test. *, p<0.05. (F) Micro-CT images of representative KRAS^LSL-G12C/+^; p53^f/f^ mice lungs at day 0, day 57, and day 150 of treatment with vehicle or MRTX849. Yellow circles depict representative shrinking tumor nodules between days 0 and 57 of treatment, and white circles outline representative growing tumor nodules between days 57 and 150. (G) Waterfall plot shows percent change in tumor area of vehicle-treated cohorts between days 0 and 57 (blue), MRTX849-treated cohorts between days 0 and 57 (red), and MRTX849-trated cohorts between days 57 and 150 (green). Representative lung H&E images shown below. (H) Micro-CT images of representative KRAS^LSL-G12D/+^; p53^wm/wm^ mice lungs at day 0, day 42, and day 153 of treatment with vehicle or MRTX1133. Yellow circles depict tracked representative shrinking tumor nodules between days 0 and 42 of treatment, and white circles outline representative growing tumor nodules between days 42 and 153 of MRTX1133 treatment. (I) Waterfall plot shows percent change in tumor area of vehicle-treated mice between days 0 and 42 (blue), MRTX1133-treated mice between days 0 and 42 (red), and MRTX1133-treated mice between days 42 and 153 (green). Representative lung H&E images shown below.

**Supplemental Figure S2.**
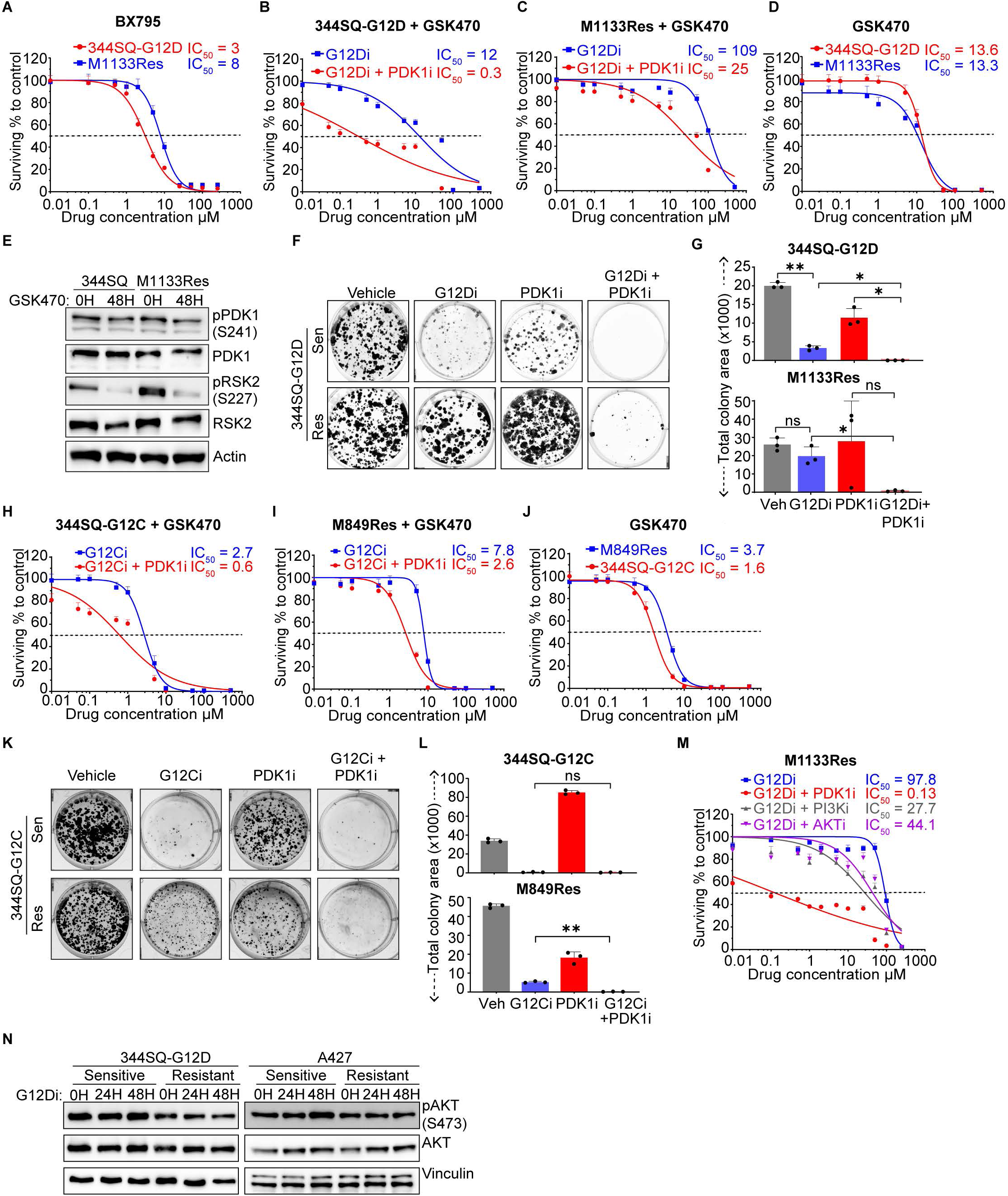
Reduction in PDK1 activity makes KRAS-mutant NSCLC more sensitive to KRAS inhibition. (A) Growth inhibition assays of 344SQ-G12D (red) and M1133Res (blue) lines when treated with BX795 at the indicated concentrations to determine IC50 values of each cell line. (B) Growth inhibition assays of 344SQ-G12D cells treated with MRTX1133 (blue) or cotreated with GSK2334470 (PDK1 inhibitor hereafter named GSK470) at its IC50 concentration (red). (C) Growth inhibition assays of M1133Res cells treated with MRTX1133 (blue) at the indicated concentrations or cotreated with GSK470 at its IC50 concentration (red). (D) Growth inhibition assays of 344SQ-G12D (red) and M1133Res (blue) cells when treated with GSK470 at the indicated concentrations. (E) Western blot analysis of 344SQ-G12D and M1133Res lines treated with GSK470 at IC50 concentration for the indicated time points to evaluate levels of phosphorylated and total PDK1 (S241) and RSK2 (S227).. (F) Colony formation assays of 344SQ-G12D and M1133Res cells treated with vehicle, MRTX1133, GSK470, or combination. (G) Quantification of the data in F. *, p<0.05. **, p<0.01. (H) Growth inhibition assays of 344SQ-G12C cells treated with MRTX849 (blue) at the indicated concentrations or cotreated with GSK470 at its IC50 concentration (red). (I) Growth inhibition assays of 344SQ-G12C M849Res cells treated with MRTX849 (blue) at the indicated concentrations or cotreated with GSK470 at its IC50 concentration (red). (J) Growth inhibition assays of 344SQ-G12C (red) and M849Res (blue) cells treated with GSK470 at the indicated concentrations. (K) Colony formation assays of 344SQ-G12C and M849Res cells treated with vehicle, MRTX849, GSK470, or combination. (L) Quantification of the data in K. **, p<0.01. (M) Growth inhibition assays of M1133Res cells treated with MRTX1133 or cotreated with either BX795 (red), PI3Ki (Buparlisib, grey), or AKTi (MK2206, purple) at their IC50 concentrations. (N) Western blot of 344SQ-G12D and A427 sensitive and resistant lines treated with MRTX1133 at IC50 concentration to evaluate total and pAKT (S473) levels.

**Supplemental Figure S3.**
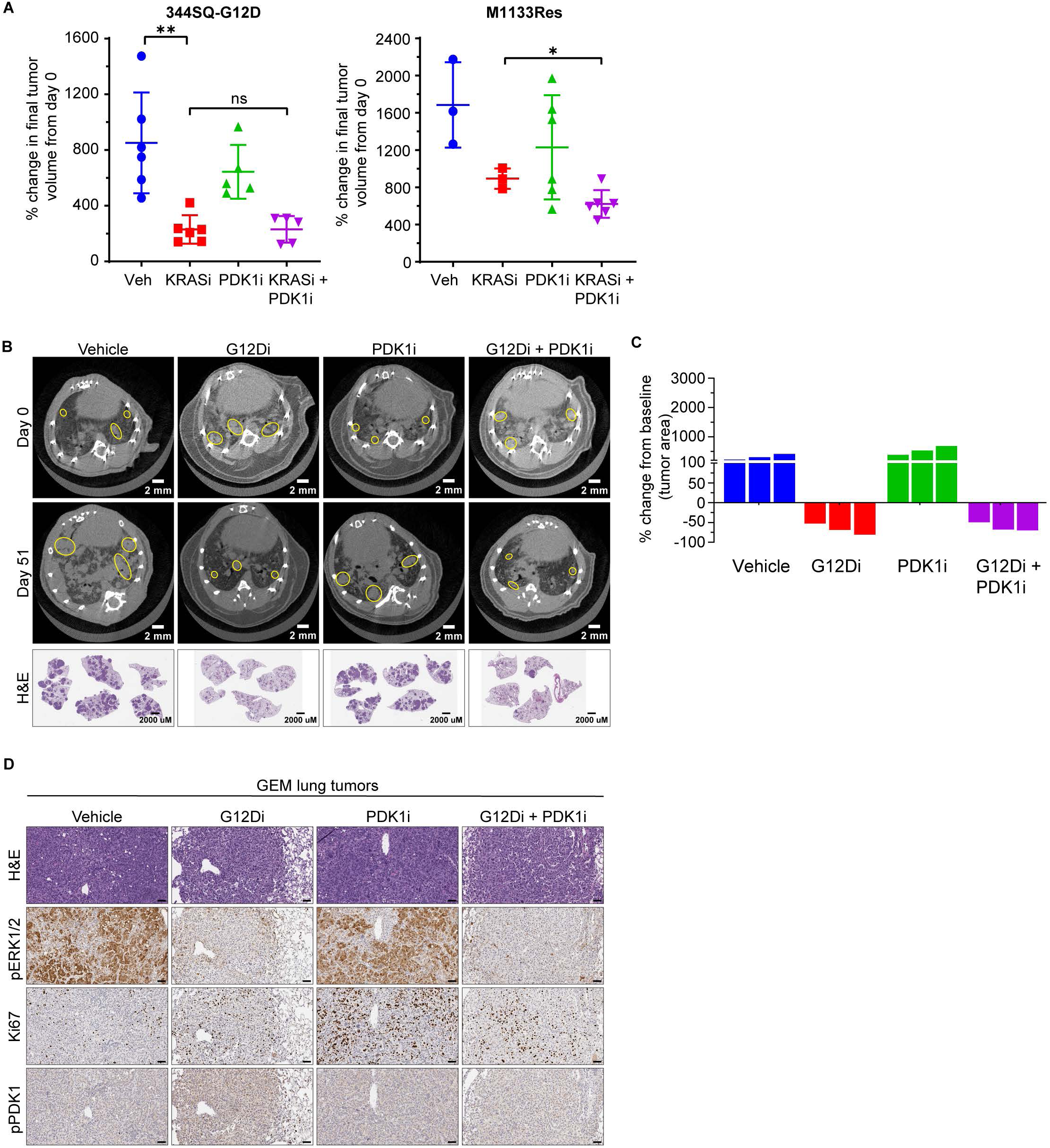
Combination treatment of resistant tumors with G12Di and PDK1i results in lower rate of tumor growth. **(A)** Percent change in tumor volumes of 344SQ-G12D (left panel) and M1133Res (right panel) cohorts from figure 3A at endpoint (day 24 and day 20, respectively). Statistical significance was calculated using Student’s t-test. *, p<0.05. **, p<0.01. **(B)** Representative micro-CT images of lungs from KRAS^G12D/+^; p53^-/-^ or KRAS^G12D/+^; ^p53R172H/R172H^ mice at day 0 and day 51 of treatment with vehicle, MRTX1133, GSK470 (PDK1i), and combination of MRTX1133 and GSK470. Day 0 images were taken 2 months after Adenoviral-Cre administration. Yellow circles outline target lesions. The bottom row depicts representative lung H&E images of mice from the respective cohorts collected at day 51. **(C)** Waterfall plot illustrates the percent difference in tumor area between day 0 and day 51 of treatment. Tumor area was calculated from three independent tumor nodules per mouse as measured from micro-CT images. **(D)** IHC staining of representative lung tumor nodules from mice treated in figure 3C. The sections were stained for pERK1/2 (T202/Y204), Ki67, and pPDK1 (S241). H&E images are shown on the top. Scale bars correspond to 50 µM.

**Supplemental Figure S4.**
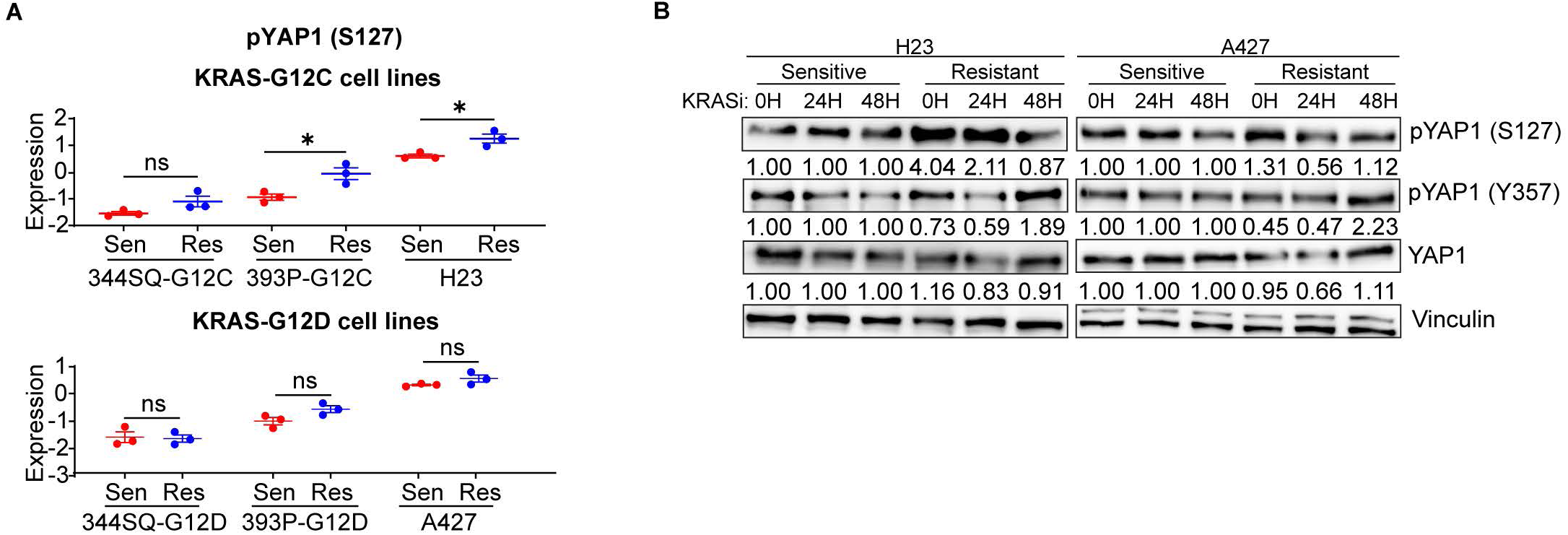
Increased YAP1 activation in KRAS^G12C^ and KRAS^G12D^-driven NSCLC with acquired resistance to KRAS inhibition. **(A)** Dot plots showing pYAP1 (S127) levels from RPPA analysis of three KRAS^G12C^ and KRAS^G12D^ sensitive and resistant lines treated with MRTX849 or MRTX1133 at respective IC50 concentrations for the indicated time points. Statistical significance was calculated using Student’s t-test. *, p<0.05. **(B)** Western blotting of H23 and A427 sensitive and resistant lines treated with MRTX849 and MRTX1133 respectively for the indicated time points. The blots were probed for pYAP1 (Y357), pYAP1(S127). and YAP1.

**Supplemental Figure S5.**
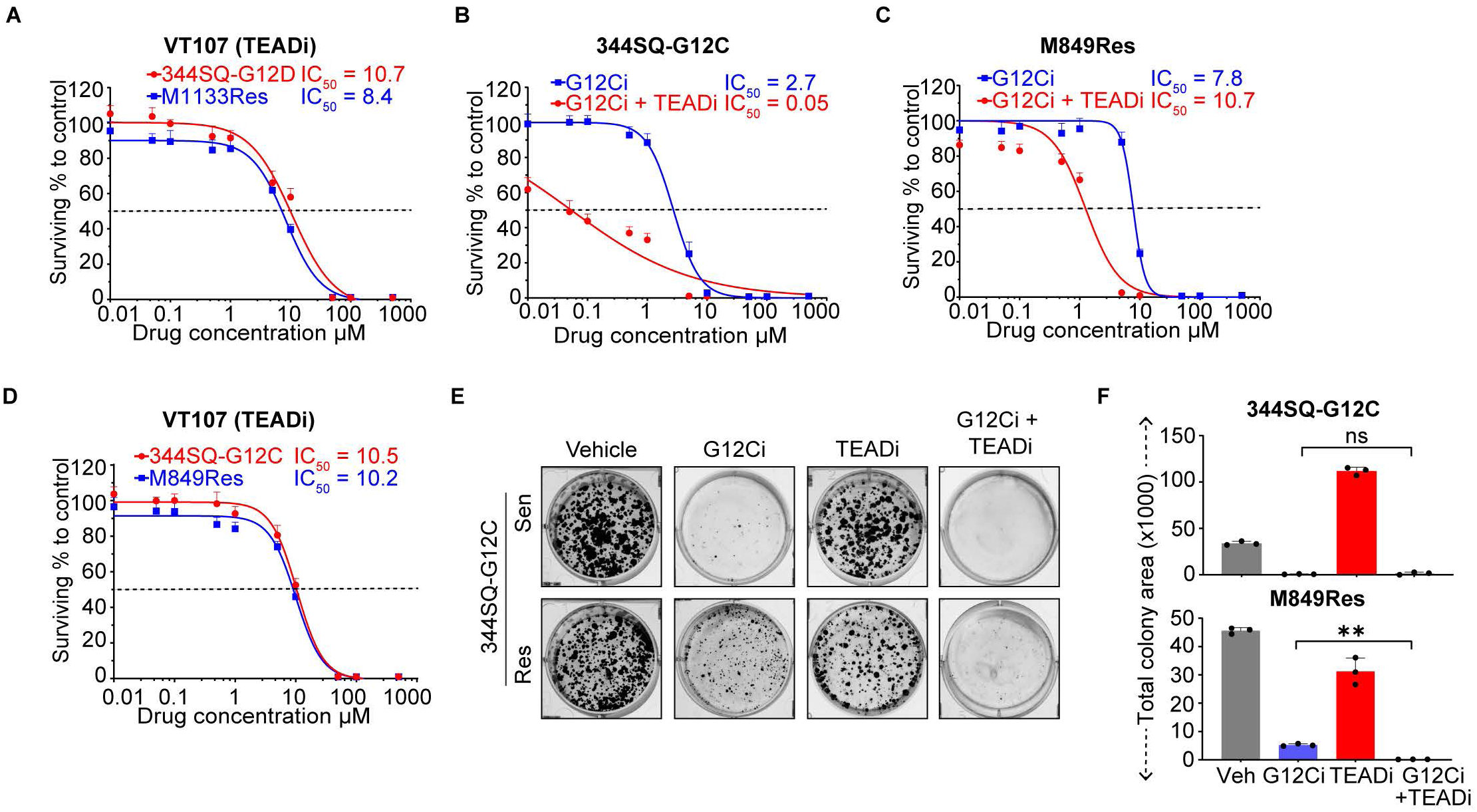
Inhibition of YAP1/TEAD activity makes KRAS-mutant NSCLC more sensitive to KRAS inhibition. **(A)** Growth inhibition assays of 344SQ-G12D (red) and M1133Res (blue) lines when treated with VT107 at the indicated concentrations for 72 hours. **(B)** Growth inhibition assays of 344SQ-G12C cells treated with MRTX849 (blue) at the indicated concentrations or cotreated with VT107 at its IC50 concentration (red) for 72 hours. **(C)** Growth inhibition assays of M849Res cells treated with MRTX849 (blue) at the indicated concentrations or cotreated with VT107 at its IC50 concentration (red) for 72 hours. **(D)** Growth inhibition assays of 344SQ-G12C (red) and M849Res (blue) cells treated with VT107 at the indicated concentrations for 72 hours. **(E)** Colony formation assays of 344SQ-G12C and M849Res cells treated with vehicle, MRTX849, VT107, or combination. **(F)** Quantification of the data in E. **, p<0.01.

**Supplemental Figure S6.**
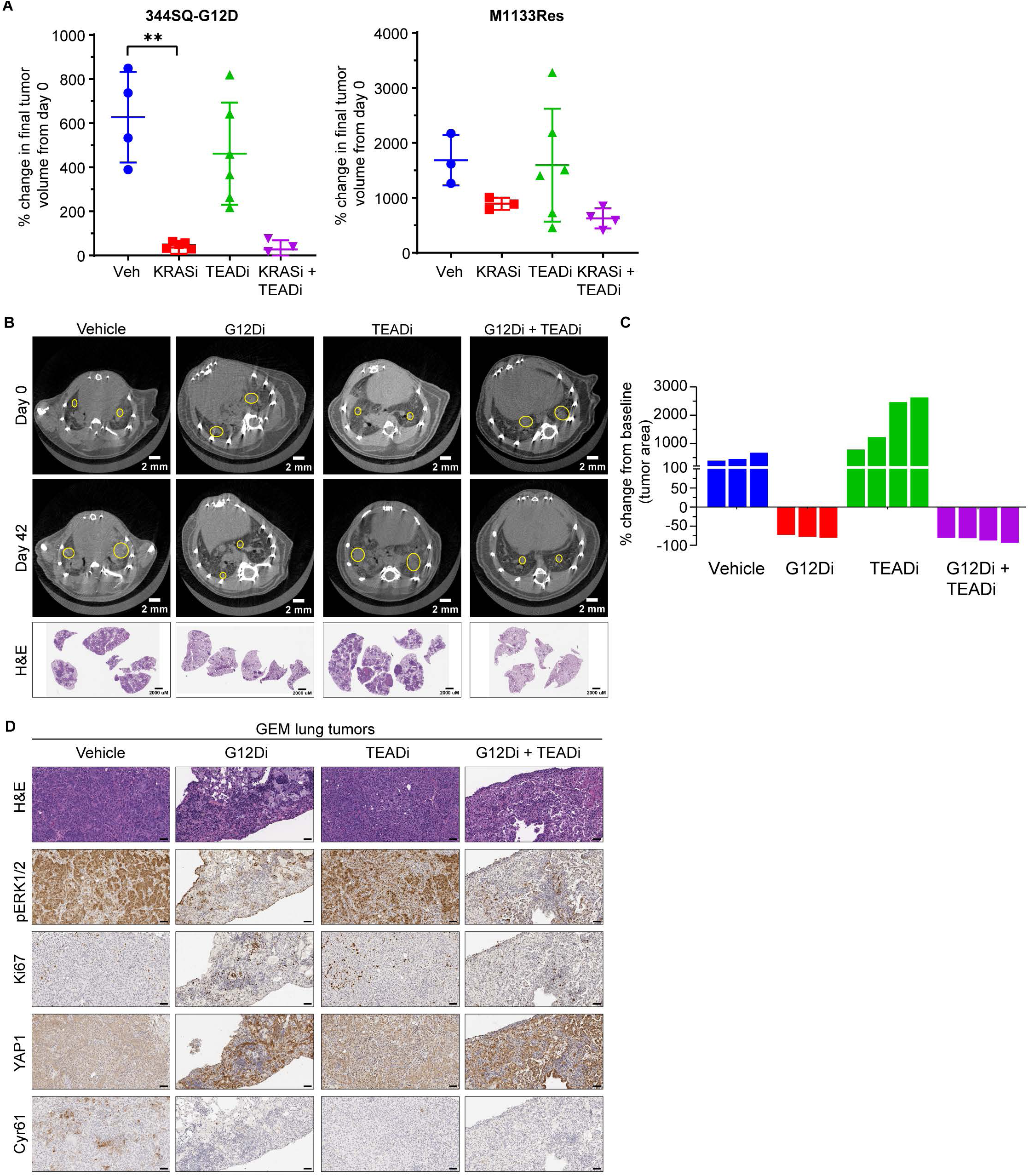
Combination treatment of resistant tumors with G12Di and YAP1/TEADi results in a lower rate of tumor growth. **(B)** Tumor volumes of 344SQ-G12D (left panel) and M1133Res (right panel) tumors from figure 6A at day 24 and day 20, respectively. Statistical significance was calculated using Student’s t-test. **, p<0.01. **(C)** Representative micro-CT images of KRAS^LSL-G12D/+^; ^p53wm/wm^ mice lungs at day 0 and day 42 of treatment with vehicle, MRTX1133, VT107, and combination of MRTX1133 and VT107. Day 0 images were taken 2 months after Adenoviral-Cre administration. Yellow circles outline target lesions. The bottom row depicts representative lung H&E images of mice from the respective cohorts harvested on day 42. **(D)** Waterfall plot shows percent difference in tumor area between day 0 and day 42 of treatment. Tumor area was calculated from three independent tumor nodules per mouse as measured by micro-CT images. IHC staining of lung tumor nodules from mice treated in figure 6C. The tumors were stained for pERK1/2 (T202/Y204), Ki67, YAP1 and Cyr61. H&E images are shown on the top. Scale bars correspond to 50 µM.

**Supplemental table 1.** List of antibodies used for western blot and IHC.

| **Antibody name** | **Catalog No.** | **WB dilution** | **IHC dilution** |
| --- | --- | --- | --- |
| pERK1/2 (Thr202/Tyr204) | CST 9101 | 1 in 1000 | N/A |
| pERK1/2 (Thr202/Y204) [IHC] | CST 4370 | N/A | 1 in 200 |
| Erk1/2 | CST 9102 | 1 in 1000 | N/A |
| pMEK1/2 (Ser217/221) | CST 9154 | 1 in 1000 | N/A |
| MEK1/2 | CST 9122 | 1 in 1000 | N/A |
| pS6 Ribosomal Protein (Ser235/236) | CST 2211 | 1 in 1000 | N/A |
| S6 Ribosomal Protein | CST 2217 | 1 in 1000 | N/A |
| pAKT (Ser473) | CST 9271 | 1 in 1000 | N/A |
| Akt (pan) | CST 4691 | 1 in 1000 | N/A |
| pPDK1 (Ser241) | CST 3061 | 1 in 1000 | 1 in 500 |
| PDK1 | CST 3062 | 1 in 1000 | N/A |
| pYAP1 (Ser127) | CST 4911 | 1 in 1000 | N/A |
| pYAP1 (Tyr357) | ab62751 | 1 in 1000 | N/A |
| YAP1 | CST 14074 | 1 in 1000 | 1 in 500 |
| Ki67 | CST 12202 | N/A | 1 in 200 |
| Cyr61 | CST 39382 | 1 in 1000 | 1 in 200 |
| pRSK2 (Ser227) | CST 3556 | 1 in 1000 | 1 in 200 |
| RSK2 | CST 5528 | 1 in 1000 | N/A |
| Vinculin | Sigma 05386 | 1 in 1000 | N/A |
| Anti-rabbit IgG, HRP-linked | CST 7074 | 1 in 2000 | N/A |
| Anti-mouse IgG, HRP-linked | CST 7076 | 1 in 2000 | N/A |

**Supplemental table 2.**
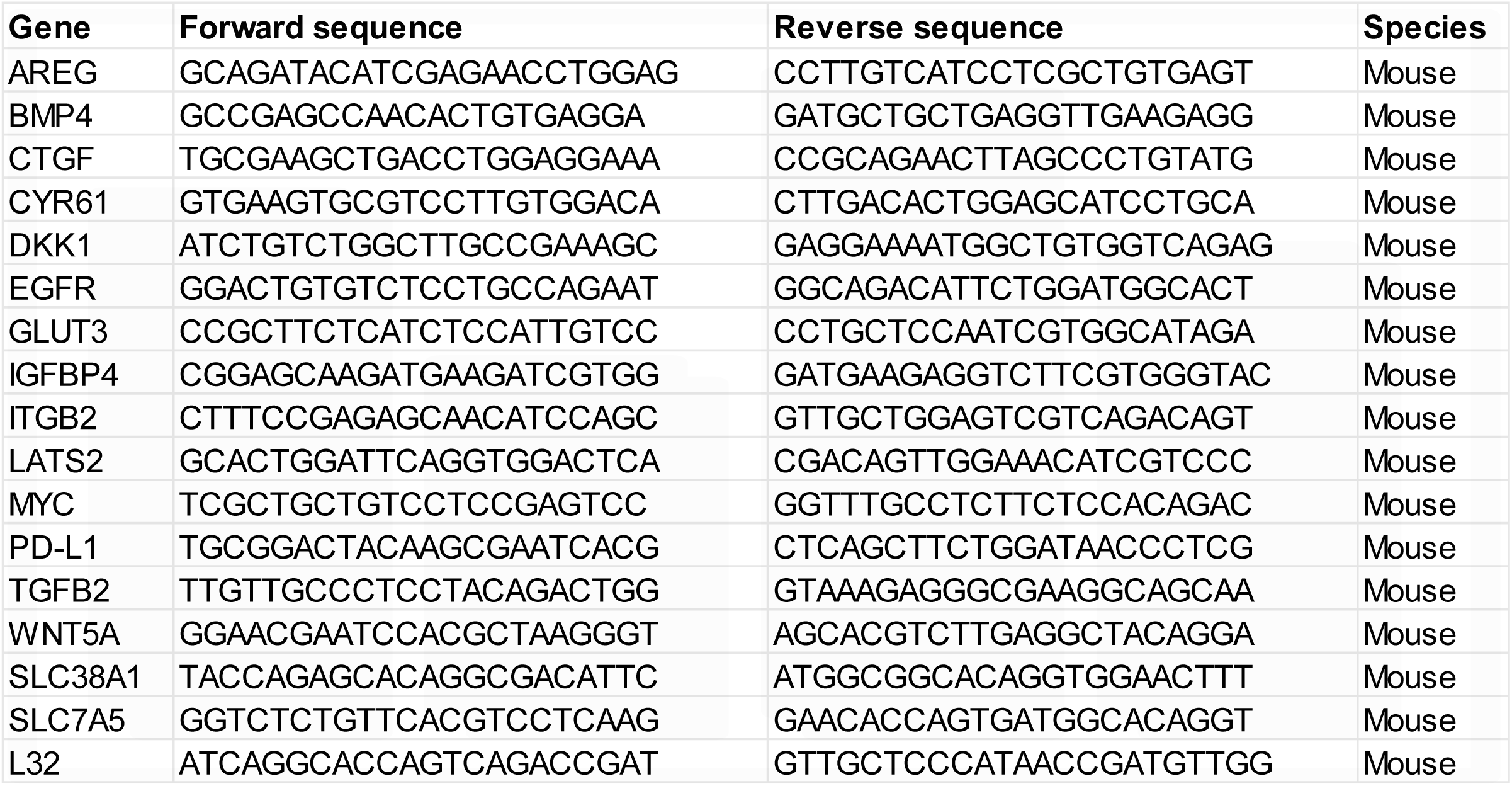
List of primers used for qRT-PCR.

